# Scalable Computational Phenomics of Nuclear Morphology

**DOI:** 10.64898/2026.09.17.752335

**Authors:** Subhadeep Duari, Raidhani Shome, Adnan Raza, Saveena Solanki, Shiva Satija, Sourav Sinha, Suvendu Kumar, Sonam Chauhan, Arushi Sharma, Vishakha Gautam, Mudit Gupta, Debarka Sengupta, Gaurav Ahuja

## Abstract

Interpretable computational pathology is constrained by a trade-off between scalable learned representations and biologically explicit phenotypes. Here we present NucXplore, a nuclear phenomics framework that converts each hematoxylin-and-eosin (H&E)-stained nucleus into 129 explicitly defined features spanning morphology, chromatin, intensity, texture, color, and spatial context. A Rust-based implementation accelerates extraction 17.6-fold over the Python reference implementation. Across three histopathology cohorts, NucXplore outperformed classical descriptors and larger deep-learning embeddings while retaining feature attribution. Applied to human skin, hierarchical multiple-instance models resolved eight cellular compartments and decoded age and sun-exposure states, revealing a largely shared aging program with age and cell-specific exposure-dependent recalibration. Conditional flow matching and neural ODE integration transformed cross-sectional phenotypes into model-implied trajectories, uncovering a midlife velocity minimum, late-life re-acceleration, and cell-specific changes in trajectory magnitude and direction. Source-free domain adaptation enabled preliminary transfer of age-associated predictions to an independent hospital cohort. NucXplore thus unifies interpretable representation, high-performance computing, and dynamical modeling for cellular phenotyping from routine histology.

## INTRODUCTION

The nucleus is a dynamic integrator of cellular state, coupling chromatin organization, nuclear envelope architecture, and mechanical forces to measurable changes in shape and internal structure. Recent structural studies have further elucidated how chromatin interacts with the nuclear lamina and how these interactions contribute to genome organization and transcriptional regulation ^1^ ^2^. Nuclear morphology, therefore, represents more than a diagnostic appearance: it provides a phenotypic readout of underlying cellular state. Consistent with this idea, nuclear features can identify cellular senescence across different stressors and tissues ^3^ and reveal age-associated changes in senescent states at single-cell resolution ^4^. Digital pathology now enables interrogation of this morphological information at an unprecedented scale. Deep learning and self-supervised foundation models trained on large histopathology collections have achieved strong performance across diagnostic, prognostic, and molecular prediction tasks ^5^ ^6^ ^7^. Yet their representations are typically encoded as hundreds to thousands of learned latent dimensions whose individual biological meanings are not explicitly defined. Classical nuclear morphometry offers the opposite advantage: direct biological interpretability, but generally captures a restricted subset of nuclear phenotype. Recent studies showing that interpretable nuclear features alone can detect senescence emphasize the information contained in morphology ^3^ ^4^, but a comprehensive representation capable of jointly resolving nuclear shape, chromatin organization, staining, texture, and tissue context at whole-slide scale remains lacking.

Human skin provides a powerful setting for addressing this problem because its aging reflects the interplay among chronological processes, cellular identity, and cumulative environmental exposure. Single-cell transcriptomics has demonstrated that human skin aging is strongly cell-type dependent, with distinct molecular programs emerging across epidermal and dermal populations ^8^. More recent studies further emphasize the heterogeneity of senescence across cell types and tissues ^9^ and implicate specific cellular compartments, including the vascular niche, in driving aspects of intrinsic skin aging ^10^. Molecular clocks can also recover chronological information from human skin, including via skin-specific DNA methylation signatures ^11^ ^12^. However, these approaches require molecular profiling and do not establish whether standard H&E histology contains an interpretable, cell-resolved record of aging and environmental exposure. More importantly, aging is not simply an endpoint to be classified, but a continuous and heterogeneous process. Single-cell studies increasingly indicate that aging involves both coordinated and stochastic changes that evolve across the lifespan ^13^. For tissue morphology, however, most computational approaches remain focused on classification or age prediction rather than reconstructing how cellular phenotypes change through age. It therefore remains unclear whether nuclear morphology can resolve continuous cell-specific aging trajectories, whether these trajectories are shared among skin compartments, and whether chronic environmental exposure alters their magnitude, timing, or direction.

Here we introduce NucXplore, an interpretable nuclear phenomics framework that represents individual H&E-stained nuclei using 129 explicitly defined features spanning morphology, chromatin, intensity, texture, color, and spatial organization. We first establish the discriminative capacity and computational scalability of this representation across three independent histopathology datasets and benchmark it against classical descriptors and deep-learning embeddings. We then apply NucXplore to human skin to resolve eight major cellular compartments and determine how their nuclear phenotypes vary with chronological age and sun exposure. Finally, we use conditional flow matching to move beyond static classification and to infer continuous, model-implied population trajectories of nuclear morphological aging, thereby resolving exposure-dependent changes in trajectory magnitude and direction across cellular compartments. Transfer to an independent hospital cohort further tests whether these nuclear aging signatures generalize beyond the source dataset. Together, NucXplore establishes interpretable nuclear phenomics as a framework for resolving cell-specific trajectories of human skin aging directly from routine histopathology.

## RESULTS

### NucXplore defines an interpretable nuclear phenotype at whole-slide scale

To construct an interpretable representation of nuclear state, we developed NucXplore, which converts each hematoxylin-and-eosin (H&E)-stained nucleus into a fixed 129-dimensional phenotypic profile organized across six complementary domains: morphology (32 features), chromatin (22), intensity (20), texture (26), color (26), and positional (3) **(Figure 1A)**. Rather than learning latent coordinates directly from images, each NucXplore dimension is explicitly defined and retains a direct relationship to a measurable nuclear property. Morphological features capture nuclear size, elongation, contour regularity, topology, and envelope complexity through conventional geometric descriptors, Hu moments, Fourier descriptors, and a nuclear envelope irregularity spectrum. Positional features describe nuclear location and local packing, whereas intensity features quantify the distribution and heterogeneity of intranuclear staining **(Supplementary Figure 1A, Supplementary Table 1)**. Texture features capture spatial chromatin organization using Gray-Level Co-occurrence Matrix(GLCM), Local Binary Patterns (LBP), and Histogram of Oriented Gradients (HOG) statistics. Color features quantify deconvolved hematoxylin and eosin signals and their relative abundance, whereas the chromatin domain resolves condensed nuclear regions through condensed-chromatin spatial metrics (CCSM) that quantify their abundance, morphology, spacing, radial localization, and internal texture **(Supplementary Figure 2A)**. Representative measurements can therefore be traced directly to nuclear properties such as area, roughness, condensed-chromatin organization, hematoxylin-to-eosin ratio, GLCM texture, and nearest-neighbor distance **(Figure 1B)**, establishing a phenotypic vocabulary in which individual dimensions remain directly interpretable.

**Figure 1.**
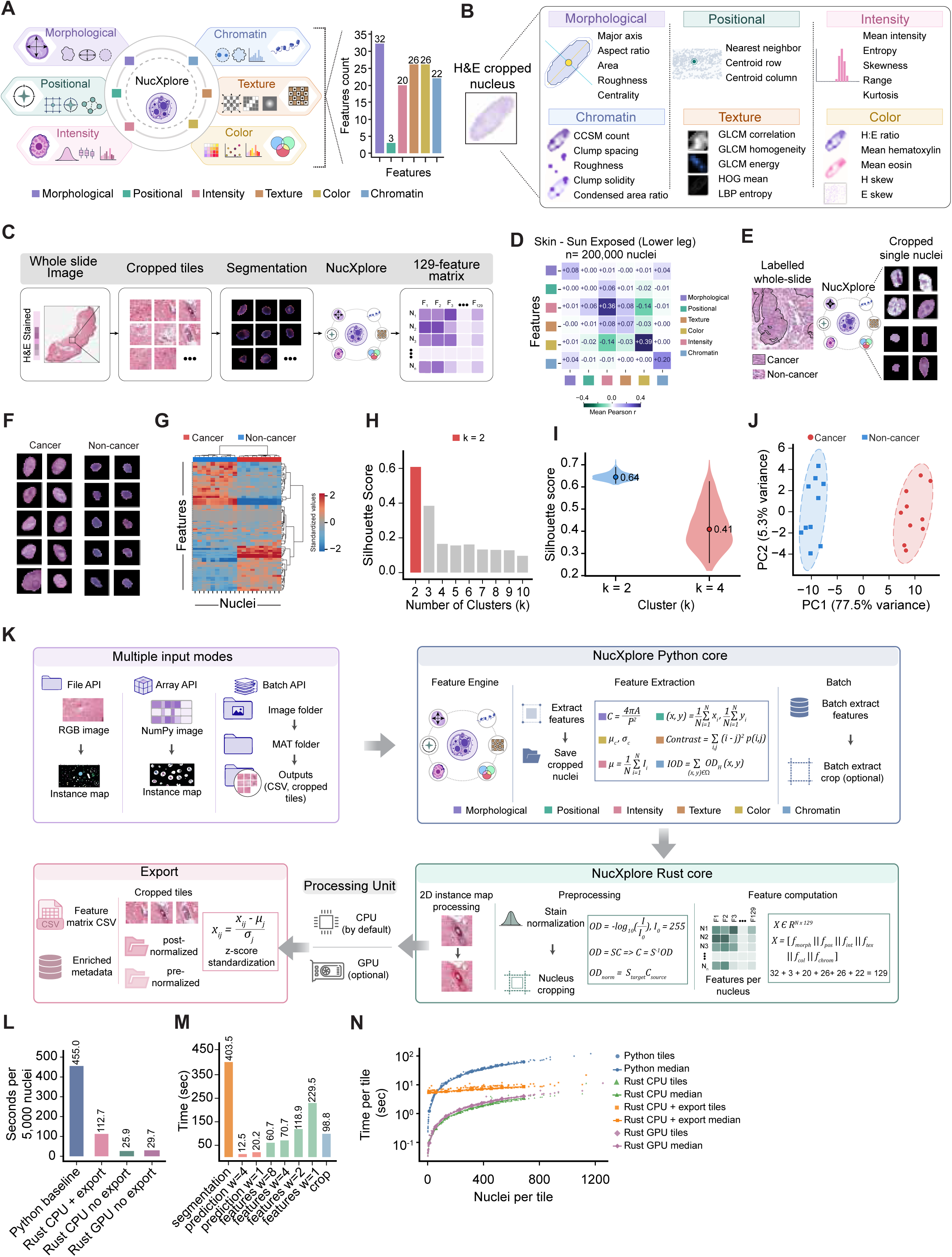
NucXplore defines an interpretable nuclear phenotype and supports high-throughput whole-slide analysis A, Overview of the NucXplore representation. Each segmented nucleus is described by 129 explicitly defined features spanning morphology (32 features), position (3), intensity (20), texture (26), color (26) and chromatin (22). **B,** Representative features mapped to a hematoxylin and eosin (H&E)-stained nuclear crop. Examples include nuclear size and contour measurements; centroid and nearest-neighbor position; intensity distribution; condensed-chromatin spatial metrics (CCSM); grey-level co-occurrence matrix (GLCM), histogram of oriented gradients (HOG) and local binary pattern (LBP) texture; and deconvolved hematoxylin and eosin measurements. **C,** Whole-slide workflow. An H&E-stained whole-slide image (WSI) is divided into tiles, nuclei are segmented and processed independently, and the resulting feature vectors are assembled into an N × 129 feature matrix. **D,** Mean pairwise Pearson correlations between the six feature domains across n = 200,000 nuclei from SE lower-leg skin. color denotes mean Pearson r (scale, -0.4 to 0.4). **E,** Application to a labelled oral squamous-cell carcinoma WSI, showing outlined cancer and non-cancer regions and representative segmented nuclei. **F,** Representative cancer and non-cancer nuclear crops, illustrating differences in contour, staining and chromatin organisation. **G,** Hierarchically clustered heat map of standardized NucXplore features across representative cancer and non-cancer nuclei. The column annotation denotes class, and color indicates the standardized feature value (scale, -2 to 2). **H,** Mean silhouette score for k-means solutions from k = 2 to k = 10, identifying k = 2 as the best-separated solution. **I,** Silhouette-score distributions for k = 2 and k = 4, with annotated scores of 0.64 and 0.41, respectively. **J,** Principal-component analysis (PCA) of standardized nuclear features. PC1 and PC2 explain 77.5% and 5.3% of the variance, respectively, and separate cancer from non-cancer nuclei; point color and shape denote class. **K,** NucXplore software architecture. File-, array- and batch-based inputs are routed through the Python interface to the native Rust engine for instance-map processing, stain normalization, nuclear cropping and feature computation. Processing uses a central processing unit (CPU) by default or an optional graphics processing unit (GPU), and outputs include feature matrices, nuclear crops and metadata. **L,** Runtime per 5,000 nuclei for the Python baseline (455.0 s), Rust CPU with crop export (112.7 s), Rust CPU without export (25.9 s) and Rust GPU without export (29.7 s). Rust CPU execution without export was approximately 17.6-fold faster than the Python baseline. **M,** Stage-resolved wall-clock times: segmentation, 403.5 s; prediction, 20.2 s with one worker and 12.5 s with four workers; feature extraction, 229.5, 118.9, 70.7 and 60.7 s with one, two, four and eight workers, respectively; and crop generation, 98.8 s. **N,** Tile-level processing time as a function of nuclei per tile for the Python, Rust CPU, Rust CPU with crop export and Rust GPU implementations. Points denote individual tiles and lines denote the corresponding medians; time is plotted on a logarithmic scale.

NucXplore was designed to preserve this nucleus-level resolution across whole histologic sections. Following image tiling and instance segmentation, each detected nucleus is processed independently and represented as a single row within an N×129 feature matrix **(Figure 1C)**. We first asked whether extending the representation across six domains simply introduced redundant measurements. Across 200,000 nuclei from sun-exposed (SE) lower-leg skin, domain-level correlation analysis showed limited coupling among most feature groups, despite stronger internal correlations within selected domains, particularly color and intensity **(Figure 1D)**. At the individual-feature level, the median Pearson correlation was only r = 0.04 for feature pairs within the same domain and approximately zero for feature pairs from different domains **(Supplementary Figure 3A)**. Domain-resolved correlation distributions further revealed heterogeneous covariance patterns within individual feature families, rather than uniform covariation across descriptors **(Supplementary Figure 3B)**. Consistent with this observation, projection of the complete feature set into a common principal-component space distributed features from all six domains across the global feature geometry rather than collapsing the representation onto a single dominant feature family **(Supplementary Figure 3E)**. Together, these analyses indicate that the six domains capture partially complementary aspects of nuclear phenotype rather than repeated measurements of a single morphological property.

We next tested whether this interpretable feature space contained sufficient intrinsic structure to distinguish biologically distinct nuclear populations without using class labels to construct the representation. In an expert-annotated oral squamous cell carcinoma section containing cancer and non-cancer regions **(Figure 1E)**, representative nuclei displayed marked differences in contour, staining, and chromatin organization **(Figure 1F)**. Hierarchical organization of the standardized 129-feature profiles produced coherent groups of nuclei that closely corresponded to the cancer and non-cancer annotations **(Figure 1G)**. An unsupervised k-means sweep identified k = 2 as the solution with the highest silhouette score**(Figure 1H)**, whereas inertia-based analysis identified an elbow at k = 4 **(Supplementary Figure 3C)**. Direct comparison of these alternatives showed substantially stronger cluster separation for k = 2 than for k = 4, with silhouette scores of 0.64 and 0.41, respectively **(Figure 1I)**. Visualization of the four-cluster solution indicated that it primarily subdivided the broader cancer-non-cancer structure rather than producing a more strongly separated partition **(Supplementary Figure 3D)**. Principal-component analysis independently recovered the same dominant biological distinction, with cancer and non-cancer nuclei occupying distinct regions along PC1, which accounted for 77.5% of the variance, compared with 5.3% for PC2 **(Figure 1J)**. Thus, biologically meaningful nuclear states emerge directly from the NucXplore representation before supervised model training.

Scaling such a representation from individual nuclei to whole-slide cohorts requires feature computation to remain tractable across populations containing millions of cells. We therefore implemented the complete NucXplore workflow through a Python-facing interface coupled to a native Rust computation engine. The architecture supports file, array, and batch-based inputs; native processing of instance maps; stain normalization; nuclear cropping; feature extraction; optional GPU execution; and export of both feature matrices and nuclear crops **(Figure 1K, Supplementary Table 2)**. In a benchmark of 5,000 nuclei, the pure-Python implementation required 455.0 s, whereas native Rust execution without crop export required 25.9 s, corresponding to an approximately 17.6-fold speedup in runtime. The GPU-enabled workflow without crop export required 29.7 s, whereas enabling crop export increased the Rust CPU runtime to 112.7 s, indicating that image-output generation accounts for a substantial portion of the remaining computational overhead **(Figure 1L)**. Stage-resolved profiling supported this interpretation. In the profiled workflow, upstream segmentation remained the largest computational component, requiring 403.5 s, whereas prediction required 20.2 s with one worker and 12.5 s with four workers. Feature extraction decreased from 229.5 s with one worker to 118.9, 70.7, and 60.7 s with two, four, and eight workers, respectively, while crop generation itself required 98.8 s **(Figure 1M)**. Independent scaling experiments across increasing worker counts and dataset sizes similarly demonstrated substantial gains from parallelization, although the incremental benefit diminished at higher worker counts **(Supplementary Figure 3F)**. At the tile level, all Rust implementations remained markedly faster than the Python reference across a broad range of nuclear densities, with processing time increasing predictably as the number of nuclei per tile increased **(Figure 1N)**. Kernel-specific CPU-GPU benchmarking further showed that acceleration was workload-dependent: GPU execution became advantageous for selected computationally intensive operations and larger image crops, particularly for HOG, whereas smaller workloads could remain CPU-favored **(Supplementary Figure 3G)**. Together, these results establish NucXplore as a compact and explicitly interpretable nuclear phenomic representation that captures complementary dimensions of nuclear biology, recovers biologically meaningful structure without supervised feature learning, and can be computed efficiently at whole-slide scale. This representation, therefore, provided the basis for testing whether interpretable nuclear phenomics could retain the discriminative capacity of substantially larger learned image embeddings across independent histopathology datasets.

### NucXplore outperforms larger representations across independent datasets

Having established that NucXplore provides a compact and biologically interpretable representation of nuclear phenotype, we next tested whether this transparency compromises discriminative performance. We benchmarked NucXplore against three conventional nuclear descriptors: morphometrics (7 features), elliptic Fourier analysis (12), and Zernike moments (25), and three substantially larger deep-learning representations: DINOv2 ViT-B/14 ^14^ (768 dimensions), EfficientNet-B0 ^15^ (1,280), and ResNet-50 ^16^ (2,048) **(Figure 2A, B)**. All representations were evaluated using the same supervised and unsupervised analysis framework across three independent H&E datasets spanning melanoma, oral squamous cell carcinoma, and lung and colon cancer **(Figure 2A)**. These cohorts differed markedly in scale and complexity, ranging from binary cancer-versus-normal classification in the Freire et al. dataset to five histologic classes in the Borkowski et al. dataset and ten nuclear classes with pronounced class imbalance in the Schuiveling et al. melanoma cohort **(Figure 2C-E)**. Despite containing only 129 dimensions, NucXplore consistently showed the strongest overall discrimination across the three cohorts **(Figure 2F-H, Supplementary Table 3-5)**. In the ten-class melanoma cohort, NucXplore achieved a mean macro-AUC-ROC of 0.702 ± 0.002, compared with 0.675 for DINOv2, 0.657 for EfficientNet-B0, and 0.652 for ResNet-50 **(Figure 2F and Supplementary Figure 5E)**. Its advantage extended across individual melanoma phenotypes, with one-versus-rest analyses showing class-dependent improvements over both conventional and learned representations **(Supplementary Figure 5D)**. In oral cancer, NucXplore reached an AUC-ROC of 0.887, exceeding DINOv2 (0.801), EfficientNet-B0 (0.791), and ResNet-50 (0.786) **(Figure 2G and Supplementary Figure 4D)**. Similarly, in the five-class lung and colon cohort, NucXplore achieved a macro-AUC-ROC of 0.953, compared with 0.941 for DINOv2, 0.937 for EfficientNet-B0, and 0.934 for ResNet-50 **(Figure 2H and Supplementary Figure 7D)**. Across accuracy, precision, recall, F1 score, Cohen’s kappa, and Matthews correlation coefficient, NucXplore remained consistently competitive with or superior to the larger learned representations **(Figure 2I; Supplementary Figure 5C; Supplementary Figure 6C; Supplementary Figure 7C)**. Thus, increasing representation dimensionality did not translate into greater discriminative power: NucXplore occupied the favorable end of the performance-dimensionality relationship in both melanoma and oral cancer while using approximately 6-fold fewer dimensions than DINOv2 and 16-fold fewer than ResNet-50 **(Supplementary Figure 5E and Supplementary Figure 6D)**.

**Figure 2.**
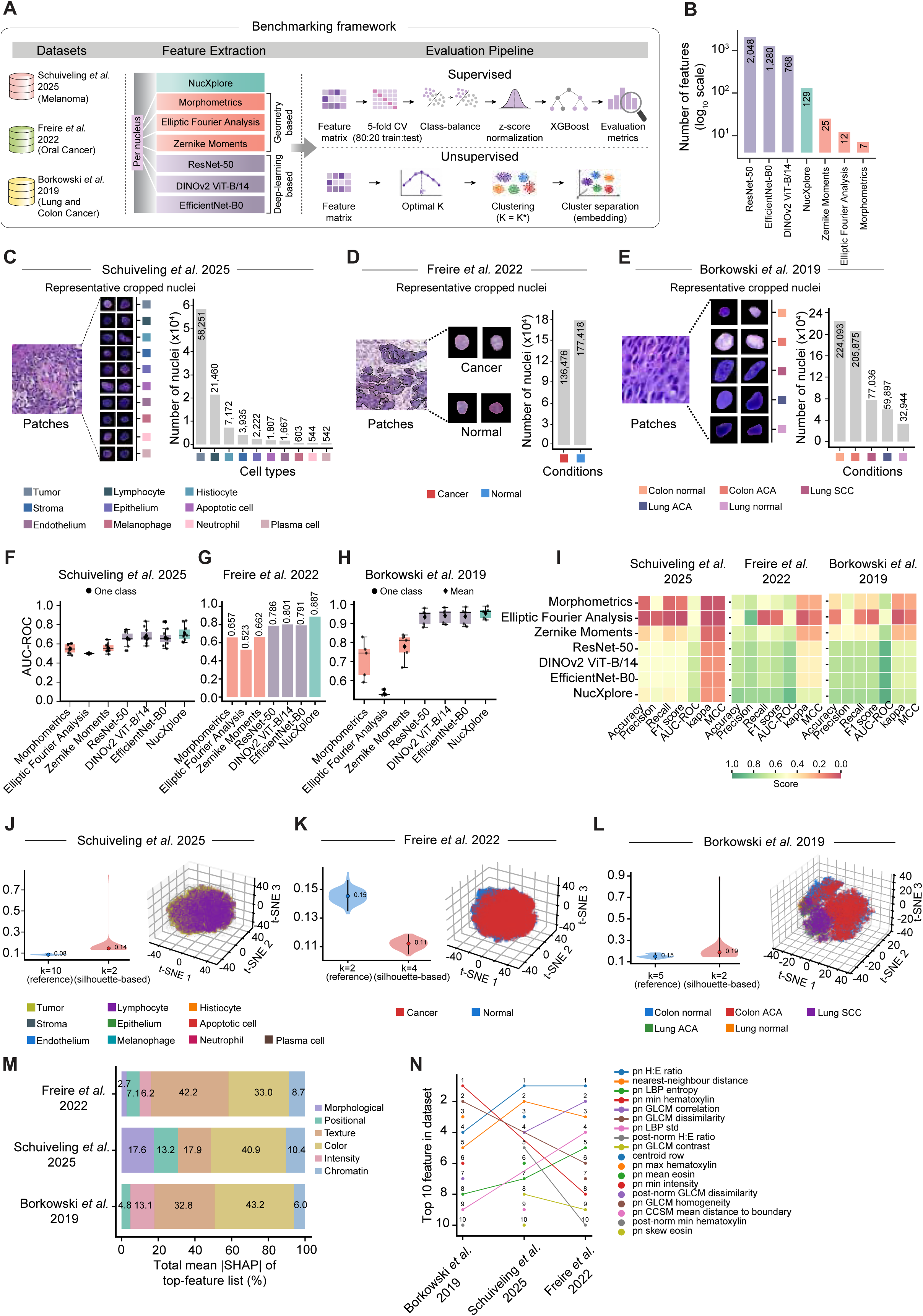
NucXplore outperforms larger nuclear representations across independent histopathology datasets A, Benchmarking design. Seven per-nucleus representations - NucXplore, morphometrics, elliptic Fourier analysis, Zernike moments, ResNet-50, DINOv2 ViT-B/14 and EfficientNet-B0 - were evaluated across PUMA melanoma, OCDC oral-cancer and LC25000 lung-and-colon cohorts. Supervised evaluation used XGBoost with five-fold stratified cross-validation, training-fold standardisation and balanced sample weights; unsupervised evaluation compared data-driven cluster solutions and low-dimensional embeddings. **B,** Dimensionality of each representation on a logarithmic scale: ResNet-50, 2,048; EfficientNet-B0, 1,280; DINOv2 ViT-B/14, 768; NucXplore, 129; Zernike moments, 25; elliptic Fourier analysis, 12; and morphometrics, 7. **C,** PUMA melanoma cohort, showing representative H&E tissue, nuclear crops and counts for tumour (58,251), lymphocyte (21,460), histiocyte (7,172), stroma (3,935), epithelium (2,222), apoptotic cell (1,807), endothelium (1,667), melanophage (603), neutrophil (544) and plasma cell (542). **D,** OCDC oral-cancer cohort, showing representative cancer and normal tissue and nuclei; 136,476 cancer and 177,418 normal nuclei were analyzed. **E,** LC25000 lung-and-colon cohort, showing representative tissue and nuclei from normal colon (224,093), colon adenocarcinoma (205,875), lung squamous-cell carcinoma (77,036), lung adenocarcinoma (59,897) and normal lung (32,944). **F-H,** Fold-level area under the receiver-operating-characteristic curve (AUC-ROC) for the seven representations in the melanoma (f), oral-cancer (g) and lung-and-colon (h) cohorts under five-fold stratified cross-validation. Box plots show the distribution across held-out folds, black circles denote individual fold values and diamonds denote means where shown; boxes span the interquartile range with a median line, and whiskers extend to 1.5 times the interquartile range. **I,** Accuracy, precision, recall, F1 score, AUC-ROC, Cohen’s κ and Matthews correlation coefficient (MCC) across representations and cohorts. color denotes metric value from 0 to 1. **J,** Melanoma (Schuiveling *et al.*, 2025). Silhouette-score distributions compare the annotated class number (*k* = 10; silhouette score = 0.08) with the silhouette-derived alternative (*k* = 2; score = 0.14). The accompanying three-dimensional t-SNE projection is coloured by the 10 annotated cell classes. **K,** Oral cancer (Freire *et al.*, 2022). Silhouette-score distributions compare the annotated class number (*k* = 2; score = 0.15) with the silhouette-derived alternative (*k* = 4; score = 0.11). The three-dimensional t-SNE projection is coloured by the annotated cancer and normal classes. **L,** Lung and colon cancer (Borkowski *et al.*, 2019). Silhouette-score distributions compare the annotated class number (*k* = 5; score = 0.15) with the silhouette-derived alternative (*k* = 2; score = 0.19). The three-dimensional t-SNE projection is coloured by the five annotated tissue and cancer classes. **M,** Relative contribution of the six NucXplore domains to total mean absolute TreeSHAP attribution among the top-ranked features in each cohort. color and texture account for the largest fractions across the three datasets, with additional contributions from morphology, position, intensity and chromatin. **N,** Rank comparison of the ten most informative NucXplore features across cohorts, showing recurrent contributions from hematoxylin-to-eosin balance, nearest-neighbor distance, LBP and GLCM texture, hematoxylin intensity and condensed-chromatin organisation.

The explicit structure of NucXplore further enabled interrogation of the basis of these predictions. Representation-specific SHAP analyses showed that conventional morphometrics concentrated predictive information in a small number of geometric variables, whereas deep-learning models distributed importance across anonymous embedding coordinates **(Supplementary Figures 4A, C, E)**. By contrast, NucXplore linked predictive importance to measurable nuclear properties. Across all three cohorts, ROC analyses independently confirmed the performance advantage of this interpretable representation **(Supplementary Figures 4B, D, F)**. When NucXplore SHAP values were aggregated by feature domains, color-derived features accounted for the largest fraction of the attribution signal across datasets, contributing approximately 33-43% of the total importance among the leading features, with additional contributions from texture, intensity, morphology, chromatin, and spatial organization **(Figure 2M)**. Importantly, the predictive signal was not restricted to staining intensity alone. Rank comparison across cohorts identified recurrent contributions from the hematoxylin-to-eosin ratio, nearest-neighbor distance, LBP and GLCM texture descriptors, hematoxylin intensity, and condensed-chromatin features **(Figure 2N)**, indicating that malignancy-associated nuclear phenotypes are encoded through coordinated changes in nuclear composition, texture, geometry, and local tissue context.

We next asked whether the same representation preserved biological organization without supervised classification. The annotated class cardinality was compared with data-driven cluster solutions in each cohort, revealing that the intrinsic organization of nuclear phenotypes did not necessarily reproduce the annotation structure one-to-one **(Figure 2J-L)**. In melanoma, the silhouette-based solution consolidated the ten annotated classes into a broader phenotypic structure **(Figure 2J)**, while repeated inertia analysis identified additional substructure with a modal elbow at k = 8, and principal-component projection revealed a dominant lower-dimensional partition **(Supplementary Figures 5A, B)**. In the binary oral-cancer cohort, the data-driven solution partitioned the annotated cancer-versus-normal structure, with an elbow at k = 4 and a corresponding separation in principal component space **(Figure 2K and Supplementary Figure 6A, B)**. The lung and colon cohort similarly showed broader phenotypic consolidation in the silhouette-based analysis, whereas the complementary inertia analysis identified an elbow at k = 4 among the five annotated classes **(Figure 2L and Supplementary Figures 7A, B)**. These differences between annotated classes and unsupervised solutions indicate that NucXplore captures phenotypic organization that can both merge morphologically related annotations and resolve heterogeneity within nominal classes rather than merely reproducing predefined labels. Collectively, these benchmarks demonstrate that explicitly interpretable nuclear phenomics does not require a trade-off in predictive performance. Across three independent cancer datasets, the 129-feature NucXplore representation matched or exceeded deep-learning embeddings containing hundreds to thousands of dimensions, while preserving direct attribution to nuclear morphology, chromatin organization, staining, texture, and spatial context. This combination of compactness, discrimination, and interpretability provided the foundation for asking whether the same nuclear phenomic representation could resolve more subtle biological variation, specifically, cellular identity and aging trajectories within human skin.

### NucXplore resolves cell-specific nuclear signatures of chronological aging and sun exposure

Having established the discriminative capacity of NucXplore across cancer cohorts, we next asked whether the same interpretable nuclear representation could resolve cellular identity and aging-related variation within normal human tissue. We applied NucXplore to H&E-stained GTEx skin sections and generated an expert pathologist-guided annotated reference set spanning eight major cellular compartments: keratinocytes, fibroblasts, hair follicles, sebaceous glands, sweat glands, arrector pili, blood vessels, and adipocytes **(Figure 3A, B)**. The resulting dataset comprised 4,479 annotated nuclei, with relatively balanced representation across classes (502-727 nuclei per class) **(Figure 3C and Supplementary Figure 8A**). Linear discriminant analysis revealed structured separation among the eight compartments, with the first three discriminant axes accounting for 44.1%, 20.3%, and 14.2% of the between-class variance, respectively **(Figure 3D)**. Unsupervised principal-component analysis showed greater overlap, consistent with partially shared nuclear phenotypes across related compartments **(Supplementary Figure 8B)**, whereas class-averaged feature profiles revealed distinct combinations of morphology, staining, texture, and chromatin features for each compartment **(Figure 3E)**. We next compared ten classifier families to determine how effectively these nuclear profiles encoded cellular identity **(Supplementary Figure 8C)**. XGBoost provided the strongest overall balance of discrimination and classification performance in the held-out comparison **(Supplementary Figure 8D)**, and five-fold cross-validation confirmed stable performance across splits **(Supplementary Figure 8E)**. One-versus-rest AUC-ROCs ranged from 0.918 for blood vessels to 0.980 for adipocytes, with all other cell types exceeding 0.96, including fibroblasts (0.973), arrector pili (0.972), sebaceous glands (0.979), hair follicles (0.969), keratinocytes (0.965), and sweat glands (0.964) **(Figure 3F, Supplementary Table 6)**. The confusion matrix showed that errors were concentrated among morphologically related compartments rather than being uniformly distributed, with recall ranging from 65.7% for blood vessels to 76.0% for keratinocytes **(Supplementary Figure 8F)**. Feature attribution remained directly interpretable: the hematoxylin-to-eosin ratio, hematoxylin and eosin statistics, GLCM texture, nuclear orientation, and nearest-neighbor distance were among the recurrent determinants of cell identity **(Supplementary Figure 8G)**. Thus, the 129-feature representation captured sufficient phenotypic structure to resolve major skin compartments without requiring learned image embeddings.

**Figure 3.**
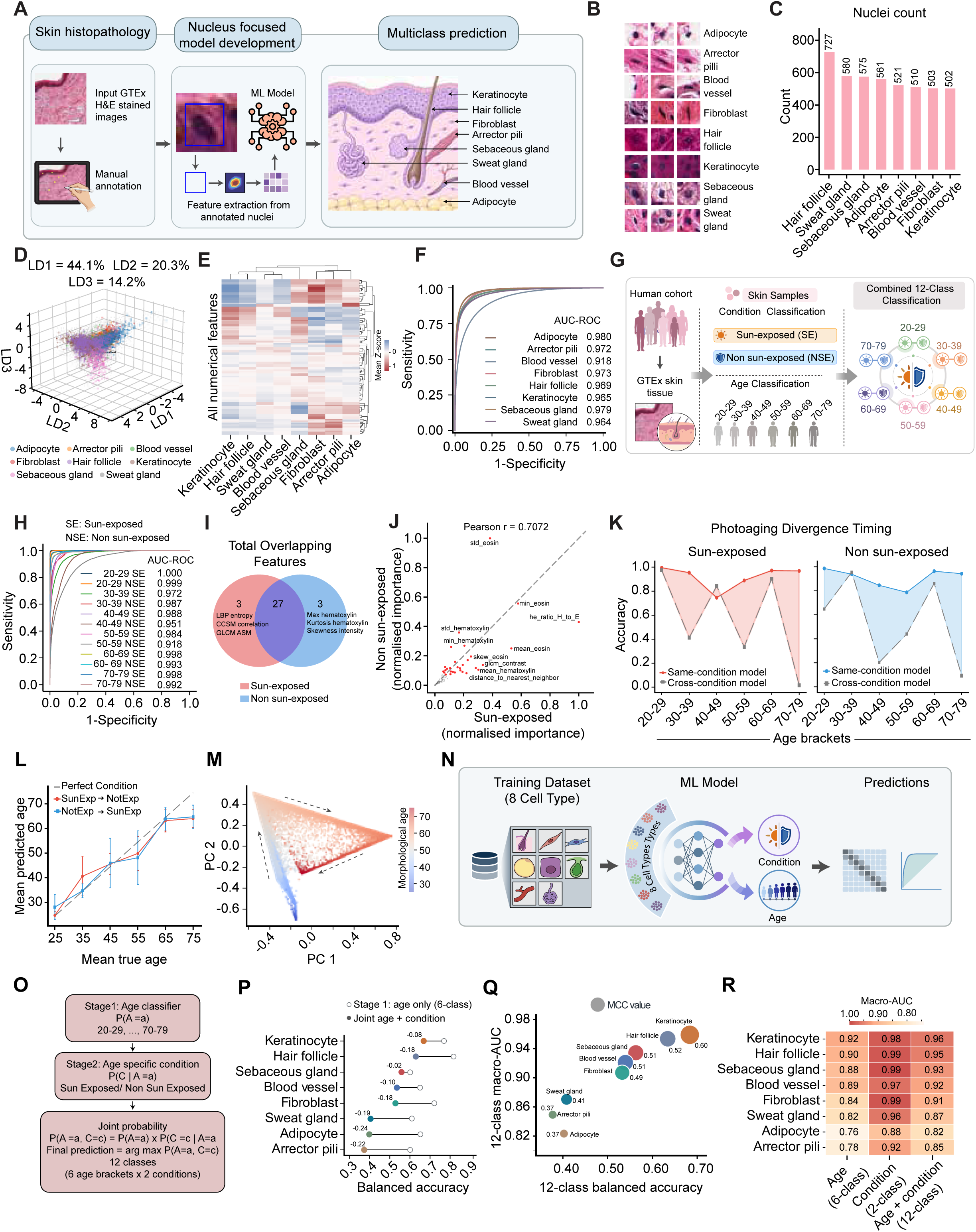
NucXplore resolves cell-specific nuclear signatures of chronological aging and sun exposure A, Workflow for generating an expert-annotated GTEx skin reference set, extracting nucleus-level NucXplore features and training a multiclass cell-type classifier. **B,** Representative annotated nuclei from adipocytes, arrector pili, blood vessels, fibroblasts, hair follicles, keratinocytes, sebaceous glands and sweat glands. **C,** Distribution of n = 4,479 annotated nuclei across the eight cellular compartments; individual classes contain 502-727 nuclei. **D,** Three-dimensional linear discriminant analysis of the annotated nuclei. LD1, LD2 and LD3 account for 44.1%, 20.3% and 14.2% of the between-class variance, respectively. **E,** Hierarchically clustered heat map of class-mean, standardized NucXplore features. color denotes the mean z-score for each cell type. **F,** One-versus-rest receiver-operating-characteristic (ROC) curves for eight-class cell-type classification; class-specific areas under the ROC curve (AUC-ROC) are shown in the panel. **G,** Hierarchical framework for classifying GTEx skin samples, represented as nucleus bags, by six age brackets and sun-exposure condition to obtain 12 combined age-condition states. SE, sun-exposed; NSE, non-sun-exposed. **H,** ROC curves for the 12 combined classes. Class-wise AUC-ROC range from 0.918 to 1.000. **I,** Overlap among the 30 most informative age-associated features from independently trained SE and NSE models. Twenty-seven features are shared, with three condition-enriched features in each model. **J,** Relationship between normalised feature importance in the SE and NSE age models. The dashed line denotes equality; Pearson r = 0.7072. Selected features are labelled. **K,** Accuracy of same-condition and cross-condition age predictions across age brackets, shown separately for SE and NSE samples. Lines connect the age-bracket estimates, and shading highlights the gap between the two prediction settings. **L,** Mean predicted age plotted against mean true age for cross-condition prediction in both directions. The dashed line denotes perfect age prediction. **M,** Principal-component projection of age-probability profiles, colored by continuous morphological age. **N,** Overview of the cell-type-specific age-condition modelling framework, in which separate models are trained for each of the eight skin compartments. **O,** Two-stage probabilistic classification. Six-class age probabilities, P(A = a), are combined with age-specific condition probabilities, P(C = c | A = a), to obtain the joint probability P(A = a, C = c) and a normalised prediction across 12 classes. **P,** Balanced accuracy of age-only and joint age-condition models for each cell type. Annotated values show joint age-condition minus age-only balanced accuracy. **Q,** Relationship between 12-class balanced accuracy and macro-AUC-ROC across cell types. Point size and labels indicate the MCC. **R**, Cell-type-specific macro-AUC-ROC for age-only, condition-only and joint age-condition classification.

We then asked whether nuclear phenotype also encoded chronological age and anatomical sun-exposure status. GTEx skin samples were organized across six age brackets from 20-29 to 70-79 years and two conditions, sun-exposed (SE) and non-sun-exposed (NSE) skin **(Figure 3G)**, with nuclei represented across all eight cellular compartments and age groups **(Supplementary Figure 8I)**. To separate the two biological signals, we used a hierarchical framework in which a six-class model first estimated age probabilities and age-specific models subsequently estimated exposure status; these probabilities were combined to generate a normalized 12-class age-by-condition prediction **(Figure 3G)**. Nuclei within each sample were encoded and aggregated using a transformer-based, gated-attention multiple-instance learning architecture with complementary bag- and nucleus-level objectives **(Supplementary Figure 8H)**. Both components were independently recoverable: age-class AUC-ROC ranged from 0.900 to 1.000, whereas condition classification reached AUC-ROC of 1.000 for SE and 0.948 for NSE skin **(Supplementary Figure 8J)**. The combined model retained strong discrimination across all 12 age-condition states, with class-wise AUC-ROCs ranging from 0.918 to 1.000 **(Figure 3H)** and an overall macro-AUC-ROC of 0.982, balanced accuracy of 0.803, and Matthews correlation coefficient of 0.745 **(Supplementary Figure 8K)**. The age-associated nuclear signal was predominantly shared between exposure conditions. Twenty-seven of the 30 most informative age features overlapped between the independently trained SE and NSE models, leaving only three condition-enriched features in each set **(Figure 3I)**. Their relative importance was also correlated across conditions (Pearson r = 0.7072; **Figure 3J**). Shared features included the hematoxylin-to-eosin ratio, eosin and hematoxylin intensity statistics, GLCM texture, and nearest-neighbor organization, consistent with a common morphological component of aging **(Supplementary Figure 8L)**. Against this shared background, differential ranking identified a smaller exposure-associated component involving stain balance, nuclear orientation, intensity dispersion, and related texture measurements **(Supplementary Figure 8M)**. These results indicate that photoaging is not represented by an entirely distinct nuclear phenotype; rather, exposure modifies the weighting and expression of a largely shared aging-associated feature program. Importantly, the effect of exposure was not constant across age. Models evaluated under the same exposure condition generally retained high accuracy, whereas cross-condition predictions exhibited pronounced losses in selected age brackets **(Figure 3K)**. The resulting age bias changed across the lifespan: cross-condition shifts were relatively small or positive in younger groups but became strongly negative in later life, reaching approximately −10 years in both prediction directions for the 70–79-year group **(Supplementary Figure 8N)**. Mean predicted-age profiles likewise showed increasing departure between same- and cross-condition predictions at older ages **(Figure 3L, Supplementary Table 15)**. Projection of the age-probability distributions produced a continuous morphological-age manifold rather than six isolated states **(Figure 3M)**, and visualization by chronological age and exposure preserved the dominant age-ordered geometry while revealing partial exposure-dependent displacement **(Supplementary Figure 8O)**. Together, these analyses show that sun exposure progressively recalibrates an underlying nuclear aging signal rather than imposing a fixed age-independent offset.

Finally, we tested whether this relationship varied among cellular compartments by training the hierarchical age-condition framework separately for each of the eight annotated populations **(Figure 3N)**. The same two-stage probabilistic structure was retained for all cell-specific models **(Figure 3O)**. Keratinocytes showed the strongest overall combined performance and only a modest reduction in balanced accuracy when moving from age-only to joint age-condition prediction, whereas adipocytes and arrector pili showed substantially larger decreases **(Figure 3P)**. Joint balanced accuracy and macro-AUC-ROC similarly positioned keratinocytes at the strongest end of the performance spectrum, with an MCC of 0.60, followed by hair follicles and sebaceous glands, whereas adipocytes and arrector pili showed lower discrimination **(Figure 3Q)**. This hierarchy was consistent across age-only, condition-only, and joint prediction, with keratinocytes reaching macro-AUC-ROC of 0.92, 0.98, and 0.96, respectively, compared with 0.76, 0.88, and 0.82 for adipocytes **(Figure 3R, Supplementary Table 7-14)**. Cell-specific 12-class ROC profiles further demonstrated substantial variation in the age-condition states that were most readily resolved within each compartment **(Supplementary Figure 9A)**, while the global 12-class confusion matrix showed that residual errors were concentrated primarily among adjacent age and exposure states **(Supplementary Figure 9B)**. Together, these results establish that interpretable nuclear morphology simultaneously encodes cellular identity, chronological aging, and sun-exposure history, but with markedly different information content across skin compartments. The predominance of shared age-associated features, coupled with age-dependent cross-condition divergence, further suggests that intrinsic aging provides a common nuclear phenotypic framework upon which photoaging imposes progressively cell-specific modifications. These observations motivated us to move beyond discrete age classification and ask how these nuclear phenotypes evolve continuously across the adult lifespan.

### Conditional flow matching reveals cell-specific nuclear aging dynamics and supports transfer across cohorts

Having shown that nuclear phenotypes encode discrete age and exposure states, we next asked whether these cross-sectional distributions could be organized into continuous, cell-specific trajectories of morphological aging. We modeled each of the eight skin compartments independently using conditional flow matching (CFM), in which a neural velocity field was trained to transport nuclear feature distributions between younger and older age states while conditioning on SE or NSE skin **(Figure 4A)**. To preserve representation across age, condition, cell type, and donor samples, nuclei were selected using a two-stage, sample-preserving FAISS (Facebook AI Similarity Search) strategy that maintained both local and global feature diversity **(Supplementary Figure 10A)**. The resulting subsets retained broad representation of age and cellular composition across both exposure conditions **(Supplementary Figures 10B, C)**, with low distributional deviation from the underlying feature space as assessed by Kolmogorov-Smirnov statistics **(Supplementary Figure 10D)**. Separate models were then trained for each cell type, with convergent flow-matching loss profiles across the eight populations **(Supplementary Figure 10E, Supplementary Table 16-18)**. Integration of the learned neural ordinary differential equations from young to old states generated distinct morphological trajectories for SE and NSE nuclei **(Figure 4B)**. These trajectories differed in curvature, orientation, and endpoint displacement across cell types, indicating that nuclear aging does not follow a single common path. To quantify exposure-dependent divergence independently of the two-dimensional visualization, identical young-reference states were propagated through the exposed and non-exposed velocity fields, and their separation was measured in standardized feature space. This separation increased progressively with modeled age in all eight cell types **(Figure 4C)**, with the largest cumulative divergence observed in fibroblasts, adipocytes, and arrector pili cells. Thus, chronic sun exposure was associated not simply with a shifted endpoint but with progressive separation of the modeled nuclear aging trajectories.

**Figure 4.**
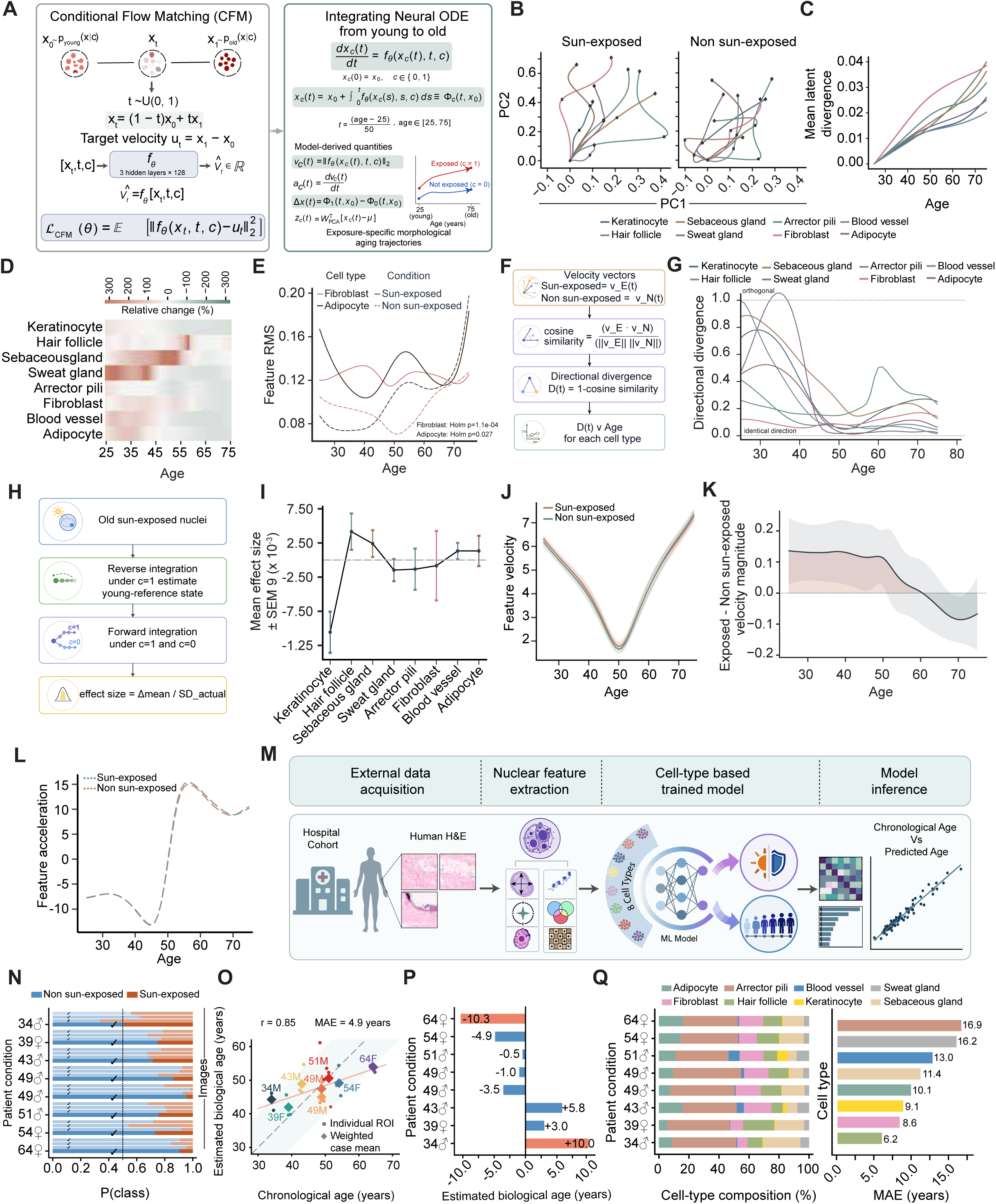
Conditional flow matching reveals cell-specific nuclear aging dynamics and supports transfer across cohorts A, Conditional flow-matching objective and integration of the learned condition-aware neural ordinary differential equation (ODE). Young and old nuclear states are connected by a probability path, and the learned velocity field is integrated over model time corresponding to ages 25-75 years. Derived quantities include feature-space velocity, acceleration, condition-dependent trajectory separation and low-dimensional phase coordinates. **B,** Qualitative PC1-PC2 phase visualisations of model-implied aging trajectories for the eight cell types under SE and NSE conditions. Separate PCA bases were fitted for the two conditions; the projections are therefore used for visualisation only and are not compared quantitatively across conditions or cell types. SE, sun-exposed; NSE, non-sun-exposed. **C,** Mean absolute trajectory separation obtained by propagating identical young reference states through the SE and NSE velocity fields and averaging the absolute paired difference across nuclei and standardized features. **D,** Relative percentage difference in the absolute value of the mean signed feature velocity across age and cell type. For each condition, values were averaged over the 20 features with the greatest temporal variance; the displayed percentage is 100 × (SE - NSE)/NSE, so positive values indicate a larger summary under the SE model. **E,** Root-mean-square (RMS) feature velocity across age for fibroblasts and adipocytes. Solid and dashed lines denote SE and NSE trajectories, respectively. Mean SE-minus-NSE differences are 0.0288 for fibroblasts and 0.0255 for adipocytes; comparisons account for autocorrelation along model age and give Holm-adjusted P = 1.06 × 10^-4^ and P = 0.0268, respectively. **F,** Calculation of directional divergence as one minus the cosine similarity between the SE and NSE velocity vectors. **G,** Directional divergence across model age for each cell type. Values of 0 and 1 denote identical and orthogonal velocity directions, respectively. **H,** Condition-switch simulation. Old SE nuclei are integrated backwards under the SE field to a model-implied young reference and then forwards under either the SE or NSE field. **I,** Mean standardized condition-switch effect size for each cell type, displayed on a ×10^-3^ scale. Points show the mean and error bars show the descriptive standard error of the mean (SEM) across retained features. These model-implied effects are derived from cross-sectional distributions and are not causal estimates of ultraviolet exposure. **J,** Equal-weight mean feature-space speed across the eight cell-type models for SE and NSE conditions. Shading denotes SEM across cell types. Both trajectories show a midlife minimum followed by late-life re-acceleration. **K,** SE-minus-NSE feature-space speed across age. Repeated-measures analysis across eight cell types gives a condition-by-age interaction of P = 0.0019 and an overall condition effect of P = 0.075. **L,** Acceleration-like numerical derivative of the condition-specific mean speed profiles with respect to normalised model time, smoothed using a five-position rolling mean. Acceleration is lower under the SE model overall (P = 0.0203), without evidence of a condition-by-age interaction. **M,** External-validation workflow comprising hospital-cohort H&E acquisition, nuclear feature extraction, cell-type assignment and age prediction; the associated source-free model-adaptation procedure is detailed in **Supplementary Figure 11A**. **N,** SE and NSE class probabilities for 24 regions of interest from eight individuals, grouped by patient. The external cohort contains 2,399,803 nuclei. **O,** Expected NSE age versus chronological age for individual regions of interest and nucleus-weighted case means. The dashed line denotes equality; across the eight case means, Pearson r = 0.85 and mean absolute error (MAE) = 4.9 years. **P,** Difference between patient-level expected NSE age and chronological age, showing overestimation in younger individuals and underestimation in the oldest participant. **Q,** Patient-level cell-type composition and cell-type-specific age-prediction MAE, which ranges from 6.2 years for hair-follicle nuclei to 16.9 years for arrector pili.

We next examined how exposure altered the magnitude of morphological change. Feature-resolved velocity differences varied markedly across age and cell type, with both positive and negative exposure-associated changes rather than a uniform acceleration of the nuclear phenotype **(Figure 4D)**. Cell-type averages likewise showed heterogeneous exposed and non-exposed velocity magnitudes **(Supplementary Figure 10F, Supplementary Table 19, 20)**. To obtain a scale-stable measure of coordinated feature movement, we calculated the root-mean-square (RMS) velocity across the feature matrix. After accounting for serial dependence along the modeled age axis and correcting across cell types, SE nuclei showed significantly greater mean RMS velocity in fibroblasts (exposed-minus-non-exposed difference, 0.0288; Holm-adjusted *P* = 1.06×10^-4^) and adipocytes (difference, 0.0255; Holm-adjusted *P* = 0.0268) **(Figure 4E)**. The remaining six cell types did not show significant differences between conditions after correction **(Supplementary Figure 10H)**, demonstrating that exposure-associated changes in overall velocity magnitude were selective rather than universal. Magnitude alone, however, does not capture whether nuclear phenotypes are changing in the same biological direction. We therefore quantified directional divergence between the exposed and non-exposed velocity vectors using one minus their cosine similarity **(Figure 4F)**. The resulting profiles were strongly age and cell-type-dependent **(Figure 4G)**. Sebaceous-gland, sweat-gland, blood-vessel, and keratinocyte models showed pronounced directional differences at younger ages, whereas hair-follicle cells exhibited a later secondary increase, and fibroblasts showed comparatively modest divergence that increased toward later life. These patterns indicate that exposure can redirect coordinated changes in nuclear features even when the overall magnitude of motion remains similar.

To further interrogate exposure-associated remodeling, we performed a condition-switch simulation. Old SE nuclei were first integrated backward under the exposed field to estimate a model-implied young reference state and then propagated forward under either the exposed or non-exposed condition **(Figure 4H)**. Standardized endpoint differences varied substantially across cell types **(Figure 4I)**. Keratinocytes showed the most negative mean shift, whereas hair-follicle nuclei showed the largest positive mean shift, with the remaining populations centered closer to zero. The complete feature-level effect distributions revealed considerably broader within-cell-type heterogeneity than these averages alone **(Supplementary Figure 10G)**. Because these simulations are derived from cross-sectional distributions rather than longitudinal interventions, they represent model-implied condition-switch effects rather than causal estimates of ultraviolet exposure. We next asked whether a common temporal structure emerged across the eight independently trained cell-type models. Equal-weight aggregation of cell-type velocity profiles revealed a pronounced U-shaped aging trajectory, with morphological velocity decreasing from early adulthood to a minimum near the 50-year model-age point before increasing again in later life **(Figure 4J)**. Importantly, the difference between exposed and non-exposed velocity changed across age rather than remaining constant **(Figure 4K).** Repeated-measures analysis across the eight cell types identified a significant condition-by-age interaction (*P* = 0.0019), whereas the overall condition effect across the complete age range was not significant (*P* = 0.075). The acceleration-like derivative showed the corresponding transition from deceleration to re-acceleration, with the strongest negative values around midlife and positive acceleration thereafter **(Figure 4L)**. In contrast to velocity, acceleration showed an overall exposure-associated reduction (*P* = 0.0203) without evidence for a condition-by-age interaction. Together, these results identify a shared midlife transition in model-derived nuclear aging kinetics while showing that sun exposure modifies its magnitude and direction in a cell-type-dependent manner. Finally, we tested whether the static age and exposure classifiers underlying NucXplore could transfer beyond GTEx to an independent hospital cohort. Routine H&E biopsies were segmented, phenotyped, assigned to cellular compartments, and analyzed using the previously trained cell-type models following source-free unsupervised SHOT (Source Hypothesis Transfer) adaptation **(Figure 4M)**. The per-sample adaptation workflow independently updated age and condition models for each cell type before weighted ensemble inference **(Supplementary Figure 11A)**. The external cohort comprised eight individuals, 24 regions of interest, and 2,399,803 nuclei. Most images were predominantly classified as non-exposed, although the inferred condition mixture varied substantially among samples **(Figure 4N)**. Transfer of the exposure signal was strongly cell-type dependent, with condition accuracy ranging from 17% for sweat-gland nuclei to 96% for hair-follicle nuclei and 100% for keratinocytes **(Supplementary Figure 11D)**. Despite this heterogeneity, patient-level expected condition age showed a strong association with chronological age (Pearson r = 0.85) with a mean absolute error of 4.9 years **(Figure 4O)**. At the individual-region level, predictions showed greater variability and an overall mean absolute error of 5.9 years **(Supplementary Figures 11B, C)**. Errors also showed evidence of compression toward the center of the training age range, with overestimation in younger individuals and underestimation in the oldest participant **(Figure 4P)**. Cellular composition varied substantially among patients, and age-prediction accuracy differed accordingly among compartments, ranging from a mean absolute error of 6.2 years for hair-follicle nuclei to 16.9 years for arrector pili cells **(Figure 4Q)**. Together, these analyses extend NucXplore from static age-associated phenotypes to a continuous, cell-resolved representation of nuclear aging dynamics. The models reveal a shared midlife slowing followed by late-life re-acceleration, while sun exposure progressively modifies both the magnitude and direction of morphological change in a cell-specific manner. Transfer to an independent clinical cohort further indicates that these nuclear aging signatures remain detectable across histologic domains, while also defining the cell-type and sample-level variability that accompanies such transfer.

## DISCUSSION

NucXplore establishes nuclear morphology as an interpretable phenotypic layer that can be quantified at scale from routine histology. This positioning is distinct from the prevailing direction in computational pathology, where increasingly large foundation models learn highly transferable representations from millions of image patches or whole-slide images ^5^ ^7,17^; Such models provide remarkable predictive power, but their representations typically comprise hundreds to thousands of latent dimensions without predefined biological meaning. NucXplore addresses a complementary problem: rather than maximizing representation complexity, it preserves the nucleus as the analytical unit and describes it through explicitly measurable properties of morphology, chromatin organization, staining, texture, and spatial context. The ability of this compact representation to match or exceed substantially larger embeddings across independent cancer datasets suggests that biological interpretability need not necessarily come at the expense of discriminative performance. Our findings extend growing evidence that nuclear architecture records cellular aging. Nuclear morphology has previously been used to identify senescent cells using deep learning ^18^, discriminate senescence using interpretable morphological features ^3^, and characterize age-associated senescent states in different tissues ^4^. NucXplore differs conceptually in that it treats aging not as a single senescence-associated phenotype but as a multidimensional nuclear state distributed across diverse cellular compartments. The same representation captured cellular identity, chronological age, and exposure-associated differences, indicating that nuclear morphology contains overlapping but separable information about several dimensions of cellular state.

This distinction is particularly important in human skin, where aging is strongly heterogeneous across cell types. Single-cell transcriptomic studies have demonstrated lineage-specific remodeling of keratinocytes, fibroblasts, and other skin populations with age ^198^, while recent spatial analyses have emphasized the strong anatomical organization of normal human skin ^20^. Molecular approaches, therefore, provide substantially greater mechanistic resolution than histology alone. However, they require specialized assays and generally quantify molecular rather than morphological state. Conversely, methylation-based skin clocks can accurately estimate chronological age ^11^ but reduce aging to a sample-level prediction. Histology-to-omics approaches can infer molecular states from tissue images ^21^, but rely on learned multimodal representations and paired molecular measurements. NucXplore occupies a different space: it asks what aspects of aging can be recovered directly from biologically defined nuclear features in standard H&E sections. This approach revealed an important organizational structure of skin aging that would be difficult to capture using bulk age predictors or opaque image embeddings. Most informative age-associated nuclear features were shared between SE and NSE, whereas their relative importance and predictive behavior changed with age and cellular compartment. This supports a model in which exposure-associated aging modifies a common underlying nuclear aging program rather than generating an entirely independent morphological state. Moreover, the strength of this signal differed markedly across compartments, with keratinocyte-associated nuclei providing a particularly informative readout of combined age and exposure. Thus, a tissue-level estimate of “skin age” can conceal substantial cellular heterogeneity in the morphological encoding of aging. A second insight was that nuclear aging was not well described by a constant linear progression. Conditional flow matching organized cross-sectional nuclear populations into continuous model-implied trajectories and revealed a midlife reduction in phenotypic velocity followed by late-life re-acceleration. Exposure altered both the magnitude and direction of these trajectories in a cell-dependent manner. These findings complement emerging evidence from molecular studies that human aging contains nonlinear temporal transitions rather than progressing at a uniform rate. Importantly, however, the trajectories inferred here describe population transport through cross-sectional feature space; they do not represent longitudinal movement of individual nuclei. Their principal value is therefore to provide a quantitative framework for comparing the geometry and kinetics of age-associated morphological change between cellular populations and exposure conditions.

Several limitations constrain the interpretation of these findings. Most importantly, GTEx SE and NSE originate from different anatomical sites. Because normal skin exhibits strong site-specific cellular and molecular organization ^20^, the observed differences should be interpreted as sun-exposure/site-associated phenotypes rather than causal effects of ultraviolet exposure. The condition-switch analysis is similarly model-based and should not be interpreted as an experimental reversal of photoaging. NucXplore also operates on two-dimensional H&E sections and therefore cannot capture three-dimensional chromatin topology. Its measurements depend on segmentation and staining quality, and the eight annotated skin categories represent broad cellular compartments rather than the finer states resolved by single-cell and spatial transcriptomics. Finally, external validation involved only eight individuals; despite encouraging age prediction after domain adaptation, this cohort establishes the feasibility of transfer rather than clinical generalizability. These limitations define clear directions for development. Matched and sun-exposed (SE) biopsies from the same individuals, longitudinal sampling, and quantitative exposure histories will be required to separate anatomical-site effects from environmental aging. Integration with spatial transcriptomics, methylation, or proteomics could further connect individual nuclear phenotypes to the molecular programs that generate them, while finer cellular annotation and explicit cell-cell interaction graphs could extend NucXplore beyond isolated nuclei toward tissue-level phenomics. Larger multi-institutional cohorts will also be necessary to quantify the effects of sex, ancestry, anatomical site, staining protocol, and disease state. Rather than replacing foundation models or molecular profiling, NucXplore therefore provides a complementary level of biological resolution: a compact, interpretable description of how individual nuclei encode cellular identity and aging within intact tissue architecture. By revealing a largely shared nuclear aging program, its cell-specific modification with exposure, and nonlinear morphological trajectories across adulthood, NucXplore positions nuclear phenomics as a scalable bridge between routine histopathology and the cellular biology of human aging.

## MATERIALS AND METHODS

### Study design and histological cohorts

NucXplore was evaluated using histological datasets selected to test three complementary aspects of the framework: (1) representation and computational scalability at the level of individual nuclei; (2) discriminative performance across independent malignant histologies; and (3) modeling of cellular identity, chronological aging, and sun-exposure-associated phenotypes in human skin. Publicly available datasets were used for framework development and benchmarking, whereas an independent institutional skin cohort was reserved for external evaluation.

### GTEx skin-aging cohort

Whole-slide hematoxylin-and-eosin (H&E)-stained sections of histologically normal human skin were obtained from the Genotype-Tissue Expression (GTEx) histology resource. Both available skin sites were included: sun-exposed (SE) skin from the lower leg and non-sun-exposed (NSE) skin from the suprapubic region. Formalin-fixed paraffin-embedded sections scanned at 20× were downloaded in SVS format together with the corresponding donor metadata. The dataset comprised 835 SE and 805 NSE skin samples. Following WSI cropping, filtering, and nuclear segmentation, approximately 69.2 million nuclei were retained from SE tissue and 83.5 million nuclei from NSE tissue. Donors were grouped into six decadal age intervals: 20-29, 30-39, 40-49, 50-59, 60-69, and 70-79 years. These data were used for cell-compartment classification, age and exposure modeling, and continuous trajectory analysis. Because GTEx SE and NSE samples originate from different anatomical sites, exposure status was treated as a combined anatomical-site/sun-exposure variable rather than as an experimentally isolated measure of ultraviolet exposure.

### Independent clinical validation cohort

An independent cohort of archival normal-skin H&E sections was obtained from the biorepository of the Rajiv Gandhi Cancer Institute and Research Centre, New Delhi. Sample collection and use were approved by the Institutional Review Board of the Rajiv Gandhi Cancer Institute and Research Centre under protocol RGCIRC/IRB-BHR/75/2021. The cohort comprised eight individuals ranging from 34 to 64 years of age, including five males aged 34, 43, 49, 49, and 51 years and three females aged 39, 54, and 64 years. Three regions or slides were analyzed per individual, yielding 24 histological images. All samples were reviewed by a consultant pathologist to confirm the absence of overt pathological abnormalities in the analyzed skin regions. The external cohort was excluded from model development. Images were processed using the HEIP segmentation and NucXplore feature-extraction workflow as the GTEx cohort before source-free domain adaptation and age/exposure inference.

### Cancer benchmark datasets

Three independently curated H&E datasets were used to evaluate the generalizability of the NucXplore representation across malignant histologies. The melanoma cohort was obtained from the Panoptic Segmentation of nUclei and tissue in advanced MelanomA (PUMA) dataset ^22^. Primary and metastatic melanoma regions of interest scanned at 40× contained nuclei annotated into ten classes: tumor, lymphocyte, plasma cell, histiocyte, melanophage, neutrophil, stromal cell, epithelium, endothelium, and apoptotic cell. The oral squamous cell carcinoma cohort was obtained from the Oral Cavity-Derived Cancer dataset ^23^. Nuclei were assigned to tumor/region-of-interest or non-region-of-interest classes according to the supplied annotations. The lung-and-colon cohort was obtained from LC25000 ^24^ and comprised five histological categories: colon adenocarcinoma (ACA), colon normal, lung adenocarcinoma, lung squamous cell carcinoma, and lung normal. Where instance-segmentation maps were supplied, individual nuclei were recovered directly from the corresponding label maps. Otherwise, nuclei were segmented using the procedure described below. Per-class nucleus counts are reported in **Figure 2C-E**.

### Nucleus segmentation, cropping and stain normalization

Nuclei were localised from per-tile instance-segmentation maps (stored as label matrices, e.g., the inst_map field of the annotation files); where instance maps were not provided, nuclei were obtained with established H&E nucleus segmentation ^25^. Each labelled instance was isolated, and a tight bounding-box crop of the nucleus and its binary mask were extracted. Morphological descriptors were computed on the binary mask, whereas intensity, texture, color, and chromatin-based descriptors were computed on the masked RGB (Red, Green, and Blue) crop. All 129 features were computed natively by the NucXplore engine, which implements every descriptor from first principles in compiled Rust rather than by wrapping external image-analysis libraries; each feature follows the standard mathematical definition cited for its family below. To reduce technical variation attributable to staining and scanning, H&E crops were normalized using a native Rust implementation of structure-preserving color normalization based on the sparse non-negative matrix factorization procedure described by Vahadane *et al.* ^26^. RGB intensities were transformed into optical-density space, hematoxylin and eosin stain vectors were estimated and L2-normalized, and normalized images were reconstructed against a fixed reference stain appearance. Appearance-dependent measurements were calculated from both the original and normalized nuclear images, whereas stain-independent shape and positional descriptors were calculated once. The same reference normalization procedure was applied consistently to datasets entering direct representation comparisons.

### NucXplore nuclear phenomic representation

Each nucleus is encoded as a 129-dimensional descriptor organised into six biologically defined domains: morphological (32 features), texture (26), color (26), chromatin (22), intensity (20) and positional (3). Below, A denotes mask area (pixel count), P the contour perimeter and I pixel intensity; sums run over the nucleus mask unless stated otherwise. Complete per-feature definitions are catalogued in **Supplementary Table 1**.

Morphological domain (32 features): Size and shape descriptors were derived from the binary mask and a fitted ellipse of matched second moments, comprising area A, perimeter P, equivalent diameter d_eq_ = √(4A/π), convex-hull area and convexity A_conv_/A; eccentricity, solidity (A/A_conv_), extent, aspect ratio (a/b), orientation, Euler number and circularity 4πA/P^2^; the seven Hu moment invariants φ₁…φ₇ computed from normalised central moments η_pq_ = μ_pq_/μ_0_^(p+q)/2+1^; the boundary-complexity terms fractal dimension D_f_ ≈ log P/log A, roughness P^2^/A and bending energy Σκ_i_^2^; five scale-invariant elliptic-Fourier shape descriptors FD_k_ = |Z_k_|/|Z_0_| from the FFT of the complex contour; and a Nuclear Envelope Irregularity Spectrum (NEIS) from a 64-angle radial profile r(θ), with NEIS = Σ_k≥6_|F_k_|^2^/Σ_k=2..5_|F_k_|^2^, spectral energy and spectral peak mode.

Positional domain (3 features): Centroid row and column (c_row_, c_col_) in whole-slide coordinates and the Euclidean distance to the nearest neighboring nucleus, NND = min_j≠i_ ‖c_i_−c_j_‖, a local measure of cell-packing density.

Intensity domain (20 features): Mean, median, standard deviation, minimum, maximum, range, interquartile range, skewness, kurtosis and Shannon entropy H = −Σ p_i_ log₂ p_i_ of the masked grey-level histogram, each computed pre- and post-normalization.

Texture domain (26 features): Grey-level co-occurrence matrices (GLCM) over 256 levels at displacement d = 1 and four orientations (0°/45°/90°/135°, angle-averaged), yielding six Haralick descriptors (contrast, dissimilarity, homogeneity, energy, ASM (angular second moment), correlation) ^27^; rotation-invariant local binary patterns (LBP) ^28^; and histogram-of-oriented-gradient (HOG) descriptors with 8 unsigned orientation bins over 180° and L2-normalization per cell ^29^. Crops were contrast-equalised by Contrast-Limited Adaptive Histogram Equalisation (CLAHE) prior to texture computation. All texture features were computed pre- and post-normalization.

Color domain (26 features): per-channel statistics (mean, minimum, maximum, standard deviation and skew) of the deconvolved hematoxylin and eosin optical-density channels, plus the hematoxylin-to-eosin ratio, computed pre- and post-normalization.

Chromatin domain (22 features): A Condensed-Chromatin Spatial Metrics (CCSM) family segments intranuclear condensed-chromatin clumps and quantifies their count, area, eccentricity, solidity, mean nearest-neighbor distance, mean distance to the nuclear boundary, condensed-area ratio and Haralick texture (contrast, correlation, energy, homogeneity), computed pre- and post-normalization.

### High-performance Rust implementation

The NucXplore feature engine was implemented in Rust and exposed to Python using PyO3 bindings compiled with maturin. Core numerical operations were implemented using low-level Rust crates, including rustfft (fast fourier transforms), nalgebra, ndarray, and image, with multithreaded CPU execution provided through Rayon. The application programming interface supports in-memory RGB arrays and instance maps, file-based processing, and batched extraction. Entry points include extract_features, extract_features_from_files, batch_extract_features, and BatchExtractor. Optional outputs include normalized and non-normalized nuclear crops, feature matrices, and image-level metadata. Selected computationally expensive kernels, including CLAHE, GLCM, and HOG operations, can optionally be executed using portable wgpu compute shaders supporting Metal, Vulkan, or DirectX 12 backends.

### Computational benchmarking

Feature-extraction throughput was benchmarked against a pure-Python reference implementation using 5,000 nuclei. Wall-clock execution time was measured for Rust CPU processing with and without image export and for GPU-enabled execution. Stage-specific profiling separately quantified segmentation, prediction, crop generation, and feature extraction. Parallel scaling was evaluated using one, two, four, and eight CPU workers. Tile-level execution time was additionally assessed as a function of the number of nuclei per tile. Kernel-specific CPU and GPU performance was measured across increasing nuclear crop sizes for CLAHE, GLCM, and HOG operations. Benchmarking was performed on a Linux-based server equipped with two AMD EPYC 7313 16-core processors, providing 32 physical CPU cores in total, 640 GiB system memory, and an NVIDIA GeForce RTX 3090 GPU with 24 GB VRAM.

### Feature redundancy and unsupervised analysis

Feature redundancy was evaluated using pairwise Pearson correlations. Correlation values were partitioned into within-domain and between-domain comparisons, and their distributions and medians were summarized. A domain-by-domain correlation matrix was additionally constructed from pairwise feature correlations.

For unsupervised analyses, numerical features were standardized to zero mean and unit variance. Hierarchical clustering was applied to standardized nuclear profiles. K-means clustering was evaluated over candidate cluster numbers from k = 2 to k = 8 using repeated class-balanced sampling. Twenty independent clustering repeats were performed using up to 10,000 nuclei per repeat, with K-means initialized five times per fit (n_init=5) and a base random seed of 42. Cluster separation was quantified using silhouette coefficients based on Euclidean distance, evaluated on subsamples of up to 3,000 nuclei, while within-cluster inertia was used for elbow analysis.

Principal component analysis was applied to standardized representations to characterize dominant variance structure. For visualization of the cancer benchmark cohorts, three-dimensional t-Distributed Stochastic Neighbor Embedding (t-SNE) was additionally applied ^30^. t-SNE was performed using PCA initialization, a perplexity of 30, an automatically determined learning rate, and a random seed of 42; the default scikit-learn iteration limit was retained. Class labels were not used to fit the unsupervised embeddings or clustering solutions and were overlaid only for interpretation.

### Benchmark representations

NucXplore was compared with six per-nucleus representations extracted from the same post-normalised crops. Three classical descriptors were used: basic morphometrics (7 features: circularity, eccentricity, solidity, elongation, compactness, convexity and extent); elliptic Fourier descriptors (harmonic order 4, 12 coefficients) ^31^; and Zernike moments (order 8, 25 coefficients) ^32^. Three deep-learning embeddings were extracted from ImageNet-pretrained or self-supervised backbones applied to each crop: ResNet-50 (2,048-dimensional global-average-pooled activations) ^16^, EfficientNet-B0 (1,280-dimensional) ^15^, and DINOv2 ViT-B/14 (768-dimensional class-token embedding) ^14^. All representations were stored as per-nucleus Parquet tables with provenance metadata that was strictly excluded from the feature matrix at classification time.

### Supervised benchmark classification

Each representation was evaluated independently with a gradient-boosted decision-tree classifier (XGBoost) ^33^ under five-fold stratified cross-validation. Training and test partitions of each cohort were combined, and all reported metrics are out-of-fold (pooled across the five held-out folds); no separate hold-out set was used. Within each fold, features were standardized with a StandardScaler fitted on the training split and applied to the validation split, and class imbalance was corrected with balanced sample weights w_i_ = n_samples_/(n_classes_·count(y_i_)), so that minority classes (e.g., plasma cells, n = 542) were weighted equally to majority classes (e.g., tumour, n ≈ 58,000). Each representation was evaluated at its native dimensionality without dimensionality reduction. Binary and multiclass tasks were analyzed according to the annotation structure of each dataset.

### Evaluation metrics and statistical analysis

Performance was quantified by accuracy, precision, recall and F1 (macro- and weighted-averaged), AUC-ROC (one-vs-rest, macro-averaged for multi-class tasks), Cohen’s κ, and Matthews correlation coefficient, computed with scikit-learn. Per-fold values were retained, and ROC curves were smoothed for display by monotone piecewise-cubic (PCHIP (piecewise cubic Hermite interpolating polynomial)) interpolation over 500 evenly spaced false-positive-rate points. To test whether NucXplore outperformed each alternative, pairwise comparisons against NucXplore were performed per metric using the Wilcoxon signed-rank test (paired across the five folds), with a two-sided Mann-Whitney U test as fallback; significance was annotated as * P < 0.05, ** P < 0.01 and *** P < 0.001 (note that with five folds the minimum achievable Mann–Whitney U P-value is ≈ 0.008). Feature attributions **(Supplementary Figure 4)** were obtained as mean absolute TreeSHAP values ^34^ from the trained XGBoost models.

### Skin-cell annotation and reference dataset

A pathologist-guided reference dataset was generated from GTEx H&E sections to train the skin-compartment classifier. Individual nuclei were annotated into eight major histological compartments: keratinocyte, fibroblast, hair follicle, sebaceous gland, sweat gland, arrector pili, blood vessel, and adipocyte. The final reference set comprised 4,479 annotated nuclei: 727 hair-follicle, 580 sweat-gland, 575 sebaceous-gland, 561 adipocyte, 521 arrector-pili, 510 blood-vessel, 503 fibroblast, and 502 keratinocyte nuclei.

### Skin-cell classification

NucXplore profiles from the annotated reference set were used to compare ten conventional machine-learning classifiers: logistic regression, decision tree, random forest, extremely randomized trees, support-vector classification, (k)-nearest neighbors, naive Bayes, multilayer perceptron, XGBoost, and gradient boosting. Model selection was based on predictive performance across accuracy, F1 score, MCC, and AUC-ROC, after which XGBoost was used for the final cell-compartment classifier. Performance was assessed using five-fold cross-validation. One-versus-rest ROC analysis and row-normalized confusion matrices were used to characterize class-specific discrimination and error structure. TreeSHAP was used to identify the NucXplore features contributing most strongly to compartment classification.

### Hierarchical age and exposure modelling

Age and exposure were modelled hierarchically rather than as a single flat classification problem. In the first stage, a six-class model estimated the probability distribution over age brackets, with *A* denoting age group and *a* ∈ {20-29, 30-39, 40-49, 50-59, 60-69, 70-79}:

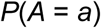

In the second stage, six independent binary classifiers estimated exposure conditional on age, with one classifier trained within each age bracket. For age group *a*, the corresponding model estimated

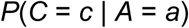

where *C* denotes exposure condition and *c* ∈ {SE, NSE. The two stages were combined probabilistically to obtain the joint age-exposure distribution:

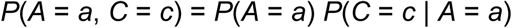

This produced 12 joint age-exposure classes. For numerical stability, the joint probabilities were normalized across all age-condition combinations before prediction:

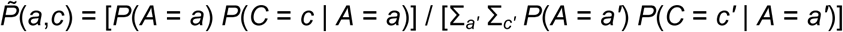

The final class was assigned as the age–condition combination with the highest normalized joint probability:

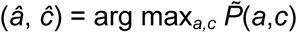

Thus, exposure prediction remained conditional on the complete Stage 1 age-probability distribution rather than being determined only from the single most probable age class.

### Multiple-instance learning architecture

Classification was performed using a transformer-based multiple-instance learning model (TransDSMIL), with the same backbone architecture used for the six-class age model and each of the six binary exposure models. Nucleus-level NucXplore feature vectors of dimension *n* were grouped into exhaustive, non-overlapping mini-bags of 256 nuclei. Each nuclear vector was first projected into a 256-dimensional latent representation using a linear projection followed by layer normalization, GELU (Gaussian Error Linear Unit) activation and dropout (0.20). The projected representations were subsequently processed by two 256-dimensional residual fully connected blocks, each containing two linear layers with layer normalization and GELU activation, followed by a final 256-dimensional projection. The encoded nuclear representations were then passed through two pre-layer-normalized transformer encoder blocks, each comprising four self-attention heads, a 512-dimensional feed-forward layer, GELU activation and dropout of 0.10.

Bag-level representations were generated using gated attention. For the latent representation *h_i_* of nucleus *i*, an unnormalized attention score was calculated as

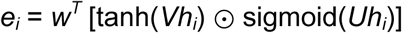

where *V* and *U* project the 256-dimensional nuclear representation into a 128-dimensional attention space, *w* projects the gated representation to a scalar score, and ⊙ denotes element-wise multiplication. Dropout of 0.25 was applied within the gated-attention module. Here *i* indexes a nucleus within a mini-bag; *h_i_* ∈ ℝ²⁵⁶ is the 256-dimensional latent representation of nucleus *i*; *e_i_* is its scalar, unnormalized gated-attention score; *V* and *U* are learned projection matrices mapping ℝ²⁵⁶ to the 128-dimensional attention space (ℝ¹²⁸); *w* is a learned 128-dimensional vector that converts the gated attention representation to a scalar; *w^T^* denotes the transpose of *w*; tanh and sigmoid are element-wise nonlinear activation functions; and ⊙ denotes the Hadamard (element-wise) product.

Attention weights were obtained by normalization across nuclei,

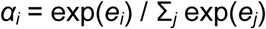

and the bag representation was calculated as the attention-weighted sum. α*_i_* is the normalized attention weight assigned to nucleus *i*, with the weights summing to 1 across all nuclei in a mini-bag; *j* is the summation index over all nuclei in that mini-bag; and *e_j_* denotes the unnormalized attention score of nucleus *j*. The softmax normalization, therefore, converts the set of scalar attention scores into a probability-like weighting over nuclei.

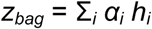

*z_bag_* ∈ ℝ²⁵⁶ is the bag-level representation obtained by summing the nucleus-level latent vectors *h_i_* after weighting each vector by its corresponding attention weight α*_i_*. It summarizes the 256 nuclei in a mini-bag as a single 256-dimensional embedding. The resulting 256-dimensional bag embedding was classified using a layer-normalized prediction head consisting of dropout (0.20), a 256-to-128 linear layer, GELU activation, a second dropout layer (0.20), and a final linear layer producing either six age-class logits in Stage 1 or two exposure-class logits in Stage 2. In parallel, an auxiliary instance-level classifier mapped each 256-dimensional nuclear representation directly to class logits. Instance-level predictions for a mini-bag were obtained by element-wise maximum pooling over nuclei:

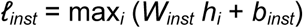

ℓ*_inst_* ∈ ℝ*^K^* denotes the pooled instance-level logit vector for the mini-bag, where *K* is the number of output classes (*K* = 6 for Stage 1 and *K* = 2 for Stage 2); *W_inst_* is the learned instance-classification weight matrix mapping the 256-dimensional latent representation to *K* class logits; *b_inst_* is the corresponding bias vector; and max*_i_* denotes element-wise maximum pooling across all nuclei *i* in the mini-bag. Thus, for each class, the instance stream retains the largest logit generated by any individual nucleus. The architecture, therefore, optimized complementary attention-weighted bag-level and maximally activated instance-level representations.

### Model training and optimization

The model was trained using a dual-stream class-weighted focal-loss objective. For a sample with true class *y*, predicted class probability *p_y_*, focal parameter *γ*, and class weight *α_y_*, the focal term was calculated from a label-smoothed target distribution. For *K* classes and smoothing coefficient *ε*, the target distribution was

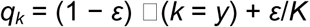

and the focal loss was

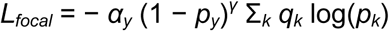

with *γ* = 2.0 and *ε* = 0.05. Class weights *α_y_* were estimated from the training-set class frequencies using balanced class weighting. In addition, training mini-bags were sampled using an inverse-frequency weighted random sampler with replacement to reduce class imbalance. The total objective combined the bag-level and auxiliary instance-level losses:

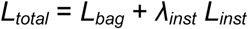

with *λ_inst_* = 0.5. Optimization used AdamW with an initial/maximum learning rate of 1 × 10^−4^, weight decay of 1 × 10^−4^, and a OneCycle learning-rate schedule. Gradients were clipped to a maximum L2 norm of 1.0, and automatic mixed-precision training was used when CUDA hardware was available. Stage 1 was trained for a maximum of 80 epochs with a batch size of 16 mini-bags and early-stopping patience of 12 epochs. Each Stage 2 age-specific binary classifier was trained for a maximum of 60 epochs with the same learning rate and batch size and an early-stopping patience of 10 epochs. In both stages, the parameter state yielding the highest validation macro-F1 score was retained for subsequent evaluation.

Before model fitting, nuclei were partitioned into training, validation and test sets in a 60:20:20 ratio using stratification over the 12 joint age-exposure labels and a random seed of 42. Feature standardization was performed using a StandardScaler fitted only to a random subsample of up to 500,000 training nuclei, after which the fitted transformation was applied to the corresponding nuclear pools. Mini-bags were generated independently within each split. Stage 1 mini-bags were pooled according to age bracket while combining the two exposure conditions, whereas Stage 2 mini-bags were formed separately for SE and NSE nuclei within each age bracket. The six Stage 2 classifiers were therefore trained independently while retaining the same TransDSMIL architecture and optimization objective as the Stage 1 model.

### Cell-type-specific age and exposure models

The hierarchical age-exposure framework was trained independently for each of the eight skin compartments to quantify the extent to which aging and exposure information differed among cellular populations. For each cell type, age-only, exposure-only, and joint age-exposure performance was quantified using balanced accuracy, macro-AUC-ROC, MCC, F1 score, precision, and recall. The same feature preprocessing and biological partitioning strategy was used across cell types.

### Representative nucleus sampling for trajectory modeling

To obtain a computationally tractable dataset while preserving biological-sample representation and nuclear phenotypic diversity, nuclei were downsampled using a sample-preserving FAISS-based procedure. Sampling was performed independently within each cell-type × age × exposure stratum. Numerical features were standardized to zero mean and unit variance before indexing. At the biological-sample level, an inverted-file flat index (FAISS IndexIVFFlat) with an IndexFlatL2 coarse quantizer was constructed using Euclidean (L2) distance. The number of IVF centroids (nlist) was determined adaptively according to dataset size, using *nlist* = 4√*N* for strata containing fewer than 10^6^ nuclei, 65,536 for 10^6^ to 10^7^ nuclei, 262,144 for 10^7^ to 10^8^ nuclei and 1,048,576 for larger datasets, with an additional constraint of approximately ≥39 observations per centroid. IVF training used up to 64*nlist* observations, subject to the available sample size, and searches were performed with nprobe=1. Within each populated IVF partition, nuclei were selected using a mixed strategy in which approximately half were chosen nearest to the corresponding centroid and the remainder were sampled randomly, thereby retaining both prototypical and within-partition phenotypic diversity. A fixed random seed of 42 was used for reproducibility. To preserve representation across biological samples, the initial patient-level FAISS selection targeted up to five times the required final sample size, with a minimum allocation of 50 nuclei per biological sample where available. The locally selected nuclei were subsequently merged and reduced to the final target by proportional sample-stratified random sampling, ensuring that each contributing biological sample retained at least one nucleus. The final representative dataset targeted 10% of the original nuclei while preserving cell-type proportions within each age × exposure stratum, with a minimum target of 100 nuclei per cell-type × age × exposure stratum. The resulting representative subset was used for conditional flow-matching analysis. Distributional similarity between the selected subset and the parent feature distributions was assessed using Kolmogorov–Smirnov statistics.

### Nuclear feature data and preprocessing for Neural ODE modelling

Nucleus-level feature tables from the representative two-level FAISS subset were analysed separately for eight skin cell types (keratinocyte, hair follicle, sebaceous gland, sweat gland, arrector pili, fibroblast, blood vessel and adipocyte) across SEand NSEtissue sites. Samples were grouped into six decadal age brackets (20-29, 30-39, 40-49, 50-59, 60-69 and 70-79 years), represented for modelling by midpoint ages *a* = 25, 35, 45, 55, 65 and 75 years and mapped to a normalized age coordinate *t_age_*:

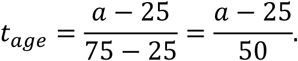

Each parquet file was treated as the sample-level unit for partitioning. The *n* features common to all eight cell-type models were standardized feature-wise within each cell type using scikit-learn StandardScaler, such that subsequent velocities and trajectory distances were expressed in standardized feature-space units:

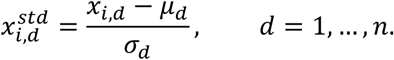

In the implemented analysis, scaling parameters were estimated from the complete per-cell-type feature matrix before sample partitioning. Unique sample identifiers were then randomly assigned to training (80%) and held-out (20%) sets (random_state = 42); the held-out samples were divided equally into feature-prediction and age-prediction subsets. No explicit age-, condition-, donor- or sex-stratification argument was supplied to the split.

### Conditional Neural ODE formulation

A separate condition-aware Neural ODE (ordinary differential equation) was fitted for each cell type *k*. Within each cell type, the standardized nuclear state *x* and continuous model time *t* were concatenated with a binary site/exposure label *c*, where *c* = 1 denoted SE tissue and *c* = 0 denoted NSE tissue. The two conditions therefore shared network parameters within a cell type, whereas parameters were not shared across cell types. The *n*-dimensional state *x_k_*(*t*) was evolved by a learned velocity field *f_θk_*:

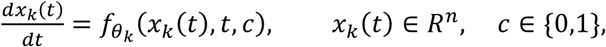

with the corresponding integral form

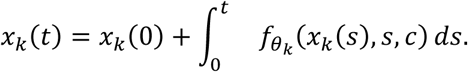

Model time was evaluated on *t* ∈ [0,1] and displayed on an age-like scale *a* = 25 + 50*t* years. Because the input data comprised cross-sectional age strata rather than longitudinal measurements, *t* was interpreted as a model-time coordinate linked to age-bin midpoints. The velocity field was implemented as a fully connected multilayer perceptron receiving the concatenated input

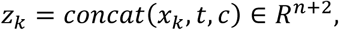

followed by three 128-unit Softplus hidden layers and a *n*-unit linear output representing *dx*/*dt*:

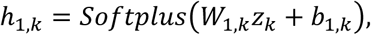

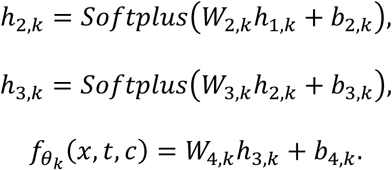

This architecture yields 257*n* + 33,408 trainable parameters per cell-type model. Linear-layer weights were initialized using Xavier-normal initialization with gain 0.1 and biases were initialized to zero.

### Conditional flow-matching objective

The velocity field was trained using conditional flow matching (CFM), without differentiating through an ODE solve during the training objective. For each training draw, *c* was sampled with probability 0.5 and young and old endpoints were independently sampled within the same condition:

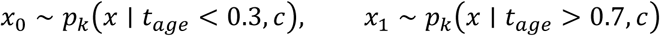

Young endpoints, therefore, corresponded to the 25- and 35-year midpoint strata, whereas old endpoints corresponded to the 65- and 75-year strata; the 45- and 55-year strata were not used as direct endpoint targets in the normal CFM training path. Endpoint pairs were not matched by donor, ROI or individual nucleus. For each pair, an interpolation coordinate τ was sampled uniformly on [0, 1], and a straight-line probability path and corresponding target velocity were constructed as

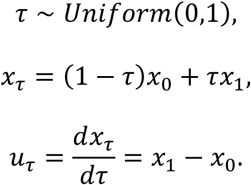

The model was trained to regress this target velocity from the interpolated state, bridge coordinate and condition by minimizing

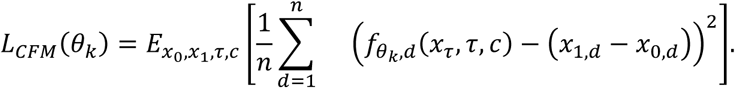

The interpolation is a training construction linking endpoint distributions and was not interpreted as the trajectory of an individual biological nucleus.

### Model training and ODE integration

For each cell type, 100,000 randomly sampled endpoint pairs were generated per epoch because *min*(100,000,10*N_train_*) reached the 100,000-pair cap was reached for all models. Models were trained for 100 epochs with mini-batches of 2,048 pairs using AdamW (initial learning rate 1 × 10^−3^; weight decay 1 × 10^−5^) and cosine annealing with *T_max_* = 100 epochs. Automatic mixed precision was used on CUDA hardware, gradients were clipped to a maximum *L*_2_ norm of 1.0, NumPy and PyTorch random seeds were set to 42, CUDA seeds were set when available, deterministic cuDNN behaviour was requested and cuDNN benchmarking was disabled. The parameter state from the epoch with the lowest mean training loss was restored after training; model selection was therefore based on training loss rather than validation-based early stopping. Across the eight training partitions, 12,256,123 image-derived nuclei were supplied to the models. Model-implied trajectories were then generated with torchdiffeq.odeint using the adaptive fifth-order Dormand-Prince solver (dopri5; relative and absolute tolerances 1 × 10^−4^), while holding *c* fixed throughout each integration. The resulting flow map is written as

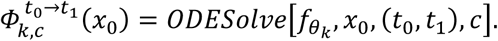

The main trajectory cache stored 51 evenly spaced evaluation positions over *t* ∈ [0,1], whereas condition-comparison and condition-switched simulations stored 50 positions; the adaptive solver could take additional internal steps. Starting nuclei for the main trajectory cache were selected separately by condition from the held-out feature-prediction subset using observed normalized age less than 0.15, which selects the 20-29-year bracket represented by age 25, and all qualifying nuclei were simulated.

### Feature-space velocity and acceleration

For each simulated nucleus *i*, cell type *k*, condition *c* and saved model-time point *t*, the fitted network returned a *n*-dimensional velocity vector

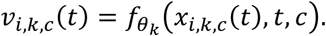

Feature-space speed was defined as the Euclidean norm of this vector,

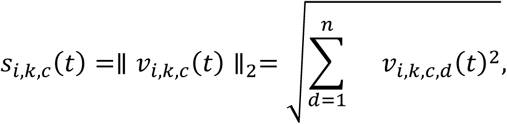

and was averaged across starting nuclei to obtain a cell-type-specific mean speed trajectory,

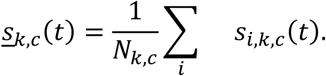

An acceleration-like quantity was calculated by numerical differentiation of the mean speed over the evenly spaced normalized-time grid,

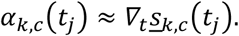

Because displayed age and model time are related by *a* = 25 + 50*t*, derivatives from the implemented analysis are expressed with respect to normalized model time; a derivative per calendar year is obtained by dividing the normalized-time derivative by 50. For the global summary, the eight cell-type mean-speed curves were averaged with equal weight at each model-age coordinate,

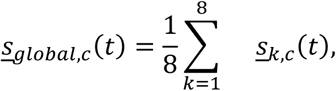

with uncertainty represented by

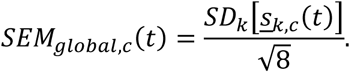

The displayed global acceleration curve was smoothed using a five-position rolling mean.

### Condition-specific trajectory separation

Condition-specific trajectory separation was evaluated within each cell type by pooling young training nuclei with normalized observed age less than 0.3 across both conditions and integrating the same starting matrix twice, once with *c* = 1 and once with *c* = 0. At each of 50 stored model-time positions, the absolute element-wise difference between the paired trajectory tensors was averaged across starting nuclei and all *n* features:

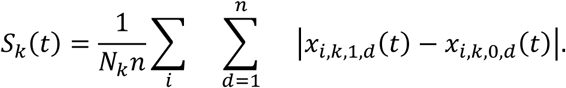

We refer to this quantity as mean absolute trajectory separation; it is distinct from dynamical-systems divergence, *▽* ⋅ *f*, which was not calculated. When an area under the separation curve was reported, the implemented summary was integrated over the displayed age coordinate in years,

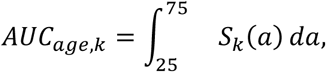

with the corresponding normalized-time quantity

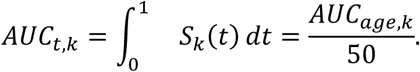

### Relative feature velocity and mean cell-type velocity

For feature-resolved analyses, signed velocities were first averaged across simulated nuclei for each feature, cell type, condition and model-time point:

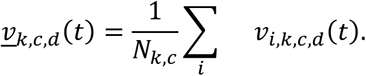

In the implemented Figure 4 analysis, features were ranked separately within each condition according to the temporal variance of their mean velocity and the 20 highest-variance features were retained,

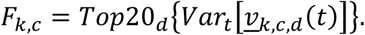

Accordingly, SE and NSE summaries could contain different feature sets. Absolute mean signed velocity was then averaged across the retained feature set,

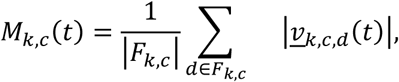

and the relative percent difference between conditions was calculated as

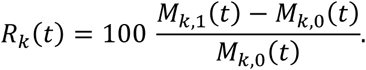

Infinite or undefined ratios produced by a zero denominator were replaced by zero in the current plotting implementation. Mean cell-type velocity was calculated independently as the arithmetic mean of the mean-speed trajectory across the 51 saved model-age positions,

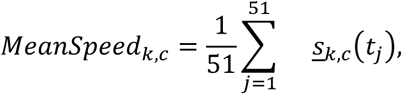

with the corresponding descriptive age-grid standard error

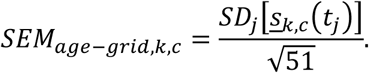

This SEM was calculated across serial age-grid positions and was therefore used only as a descriptive measure of variation along the simulated curve, not as donor-level uncertainty.

### Condition-switched feature effects

Condition-switched feature effects were estimated from old SEnuclei with normalized observed age greater than 0.7, corresponding to the 65- and 75-year midpoint strata. Each old SE state was integrated backwards from *t* = 1 to *t* = 0 under *c* = 1 to obtain a model-implied young state, which was subsequently propagated forward to *t* = 1 under *c* = 1 for reconstruction and under *c* = 0 for condition switching:

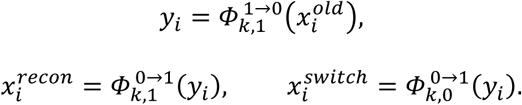

For each feature *d*, the condition-switched shift was defined as

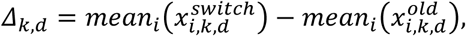

and a standardized effect size was obtained as

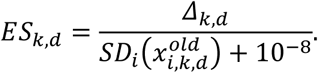

Feature-level effect-size distributions were displayed after excluding values outside *Q*_1_ − 1.5 × *IQR* and *Q*_3_ + 1.5 × *IQR* within each cell type. Means and SEM across retained features were interpreted descriptively because the *n* features are correlated measurements from the same nuclei rather than independent biological replicates.

### Low-dimensional phase trajectories

For two-dimensional visualization, each trajectory tensor of dimensions [*n_steps_*, *N*, *n*] was flattened across time and nuclei,

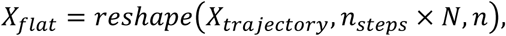

projected onto two principal components,

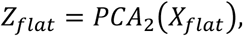

and reshaped to [*n_steps_*, *N*, 2]. Scores were averaged across starting nuclei to obtain one mean PC1-PC2 coordinate per model-age position,

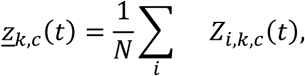

and each mean trajectory was translated so that its age-25 coordinate was at the origin,

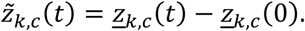

In the current phase_diagram_data.csv export, PCA was fitted separately for SEand NSEtrajectories; the two condition-specific paths are therefore represented in different coordinate bases and were not considered quantitatively comparable. The code-verified workflow additionally provides shared-PCA trajectory exports in which a single PCA basis is fitted to pooled real data within each cell type and both conditions are transformed using that basis; this shared basis should be used for the final between-condition visualization. Principal-component coordinates remain non-comparable across different cell types unless a single global PCA basis is fitted across all cell types.

### Ethics statement

Use of the independent Rajiv Gandhi Cancer Institute and Research Centre cohort was approved by the Institutional Review Board of the Rajiv Gandhi Cancer Institute and Research Centre under protocol RGCIRC/IRB-BHR/75/2021. Public GTEx and benchmark datasets were analyzed in accordance with their respective data-use conditions.

### Statistical Analysis

Statistical analyses were performed in Python, with two-sided P < 0.05 considered statistically significant unless otherwise specified. Predictive performance was quantified using accuracy, precision, recall, F1 score, area under the receiver-operating-characteristic curve (AUC-ROC), Cohen’s κ, and Matthews correlation coefficient (MCC), with macro- and weighted-averaged metrics used where appropriate for multiclass tasks. For comparisons of NucXplore with alternative nuclear representations, metric values obtained across five-fold stratified cross-validation were compared using paired Wilcoxon signed-rank tests, with two-sided Mann-Whitney U tests used as a fallback when required. Statistical significance was denoted as P < 0.05, P < 0.01, and P < 0.001. Pearson correlation coefficients were used to quantify feature redundancy and concordance of feature-importance profiles between SE and NSE skin. Distributional similarity between representative subsets selected for trajectory modelling and their corresponding parent feature distributions was evaluated using Kolmogorov-Smirnov statistics. For conditional flow-matching analyses, exposure-associated differences in aggregate nuclear phenotypic velocity were evaluated while accounting for serial dependence across the modeled age trajectory, and multiple comparisons across the eight cell types were controlled using the Holm correction, with adjusted P < 0.05 considered significant. Global age-dependent velocity and acceleration profiles were evaluated using repeated-measures analyses to assess overall exposure effects and condition-by-age interactions. Condition-switched feature effects were summarized using standardized effect sizes; values outside 1.5 × the interquartile range were excluded only from feature-level effect-distribution visualizations. Means and standard errors calculated across correlated features or serial model-age positions were treated as descriptive summaries rather than estimates of independent biological-replicate uncertainty. Associations between predicted and chronological age in the independent validation cohort were quantified using Pearson’s correlation coefficient, and prediction error was summarized using mean absolute error (MAE).

## Data availability

The data processed by the NucXplore package is publicly available through the GitHub repository (https://github.com/the-ahuja-lab/NucXplore). The data supporting the findings of this study are available in the article and its Supplementary Information, including **Supplementary Tables 1-20**, which provide the underlying feature definitions, benchmarking results, classification performance, cross-condition analyses, conditional flow-matching results, and external validation summaries. The histopathology data from the independent external-validation cohort are not publicly available but may be made available upon request. Publicly available histopathology datasets used in this study, including GTEx, PUMA, OCDC, and LC25000, are available from their respective original repositories and sources as cited in the manuscript.

## Code availability

NucXplore feature extraction was performed entirely using a purpose-built Rust engine (Rust + PyO3, built with maturin) that implements all 129 features natively and depends only on low-level numerical and image-processing crates, including *rustfft*, *nalgebra*, *ndarray* and *image*, with Rayon used for CPU parallelism and optional GPU acceleration. The source code is released as the open-source NucXplore package and is publicly available through the project GitHub repository (https://github.com/the-ahuja-lab/NucXplore). A Python-installable distribution is available through PyPI Test (https://test.pypi.org/project/nucxplore/), and detailed documentation, usage instructions and feature descriptions are provided in the NucXplore online documentation (https://the-ahuja-lab.github.io/NucXplore/Home.html). A containerized implementation of the NucXplore cell-type prediction workflow is additionally available through Docker Hub (https://hub.docker.com/r/ahujalab/nucxplore-cell-type-prediction).

## Declaration of Interests

The authors declare no competing interests.

## Acknowledgments

The authors thank the IT-HelpDesk team at Indraprastha Institute of Information Technology Delhi (IIIT-Delhi) for assistance with computational resources, the members of the Ahuja Lab for discussions and intellectual contributions throughout the development of this work, and the Genotype-Tissue Expression (GTEx) Consortium for making the histopathology data used in this study publicly available. G.A. acknowledges support from the Ramalingaswami Re-entry Fellowship (BT/HRD/35/02/2006) and a research grant (BT/PR52020/AI/133/180/2024) from the Department of Biotechnology, Ministry of Science and Technology, Government of India, as well as an intramural Start-up Grant from Indraprastha Institute of Information Technology Delhi (IIIT-Delhi).

## Author Contributions

G.A. conceived and supervised the study. S.D. led the development of the computational framework and nuclear feature representation and performed the statistical analyses. A.R. developed the NucXplore software package. S.D. and S.C. performed manual cell annotations. S.D. and V.G. developed and implemented the GTEx histopathology image-processing pipeline. Sa.S., Sh.S., S.K., and A.S. contributed to codebase validation, testing, and quality control. S.D., R.S., M.G., and So.S. performed data analysis and visualization. D.S. contributed to the development and refinement of the model architecture. G.A. and S.D. conceptualized the figures, which were prepared by R.S. and S.D. S.D. and G.A. wrote the manuscript. All authors reviewed and approved the final manuscript.

## SUPPLEMENTARY FIGURE LEGENDS

**Supplementary Figure 1: Mathematical definitions and biological interpretation of NucXplore morphology, position and intensity features**

**A,** Feature catalogue showing the domain, software feature names, mathematical definitions and biological interpretation of nuclear size, regularity, axes and topology, Hu moments, advanced contour descriptors, contour Fourier descriptors, the nuclear envelope irregularity spectrum (NEIS), centroid and nearest-neighbor position, and first- and higher-order intensity statistics. Appearance-dependent intensity features are listed for measurements before and after stain normalization. IQR, interquartile range.

**Supplementary Figure 2: Mathematical definitions and biological interpretation of NucXplore texture, color and chromatin features**

**A,** Feature catalogue showing software names, mathematical definitions and biological interpretation for GLCM, LBP and HOG texture descriptors; deconvolved hematoxylin, eosin and H\:E color measurements; and CCSM measurements of condensed-chromatin abundance, clump morphology, topology and intraclump texture. Appearance-dependent features are listed for measurements before and after stain normalization. GLCM, grey-level co-occurrence matrix; LBP, local binary pattern; HOG, histogram of oriented gradients; H\:E, hematoxylin-to-eosin ratio; CCSM, condensed-chromatin spatial metrics.

**Supplementary Figure 3: Feature non-redundancy, unsupervised structure and computational scaling of NucXplore**

**A,** Distribution of pairwise Pearson correlations for features within the same domain and between different domains. Median correlations are 0.04 and approximately 0.00, respectively. **B,** Domain-resolved Pearson-correlation distributions for morphology, intensity, texture, color and chromatin features. The vertical reference denotes r = 0. **C,** Within-cluster inertia for k = 2-10. The inertia elbow occurs at k = 4; k = 2 marks the silhouette-optimal solution used in the main analysis. **D,** Principal-component projection of the four-cluster solution, with cluster identity indicated by color and cancer or non-cancer annotation indicated by point shape. PC1 and PC2 explain 77.5% and 5.3% of the variance, respectively. **E,** Projection of individual features into a common principal-component space, with members of each of the six domains highlighted in turn. PC1 and PC2 explain 20.3% and 13.5% of feature-level variance, respectively. **F,** Wall-clock time as a function of worker number for batches containing 32, 128, 512 or 2,048 nuclei, showing diminishing gains at higher worker counts. **G,** CPU and GPU execution times across nuclear crop sizes for contrast-limited adaptive histogram equalisation (CLAHE), GLCM and HOG kernels. Time is reported in microseconds × 10³.

**Supplementary Figure 4: Feature attributions and ROC analysis across the three benchmark cohorts**

**A,** Mean absolute TreeSHAP values for features from the six comparator representations in the PUMA melanoma cohort (Schuiveling *et al.*, 2025). Morphometric features retain explicit geometric or phenotypic meaning, whereas features derived from deep-learning representations correspond to learned embedding dimensions. **B,** Cross-validated AUC-ROC values and corresponding ROC curves for melanoma classification. For this multiclass cohort, ROC curves represent macro-averaged one-versus-rest performance. Morphometrics, elliptic Fourier analysis, Zernike moments, ResNet-50, DINOv2 ViT-B/14, EfficientNet-B0 and NucXplore are compared with a random-classifier reference; NucXplore achieves an AUC-ROC of 0.701. **C,** Mean absolute TreeSHAP values for the corresponding six comparator representations in the OCDC oral-cancer cohort (Freire *et al.*, 2022), showing the relative contribution of individual morphometric variables or learned embedding dimensions to model predictions. **D,** Cross-validated AUC-ROC values and ROC curves for binary oral-cancer classification across the seven representations. NucXplore achieves an AUC-ROC of 0.885. **E,** Mean absolute TreeSHAP values for the six comparator representations in the LC25000 lung-and-colon cancer cohort (Borkowski *et al.*, 2019). As above, morphometric variables are directly interpretable, whereas deep-model features represent learned embedding coordinates. **F,** Cross-validated AUC-ROC values and macro-averaged one-versus-rest ROC curves for multiclass lung-and-colon tissue classification. The same seven representations are compared against a random-classifier reference, with NucXplore achieving an AUC-ROC of 0.953.

**Supplementary Figure 5: Unsupervised structure and supervised performance in the PUMA melanoma cohort**

**A,** Median within-cluster inertia across repeated analyses for k = 2-20; shading shows the 95% interval. The modal elbow occurs at k = 8, whereas the annotated cohort contains ten classes. **B,** Principal-component projection of the broader two-cluster NucXplore solution. PC1 and PC2 explain 21.2% and 14.5% of the variance, respectively; black crosses denote cluster centres. **C,** Fold-wise accuracy, Cohen’s κ, MCC, weighted F1 score, weighted precision and weighted recall for the seven representations under five-fold stratified cross-validation. **D,** One-versus-rest ROC curves for apoptotic cells, endothelium, stroma, tumour, epithelium, histiocytes, lymphocytes, melanophages, neutrophils and plasma cells. Insets report the AUC-ROC for each representation, and the diagonal denotes random classification. **E,** Mean macro-AUC-ROC versus feature dimensionality. NucXplore achieves a mean macro-AUC-ROC of 0.702 using 129 features.

**Supplementary Figure 6: Unsupervised structure and supervised performance in the OCDC oral-cancer cohort**

**A,** Median within-cluster inertia across repeated analyses for k = 2-8; shading shows the 95% interval. The modal elbow occurs at k = 4, compared with two annotated classes. **B,** Principal-component projection of the four-cluster solution. PC1 and PC2 explain 23.1% and 12.3% of the variance, respectively; black crosses denote cluster centres. **C,** Fold-wise AUC-ROC, F1 score, recall, precision, accuracy, Cohen’s κ and MCC for the seven representations under five-fold stratified cross-validation. **D,** Mean AUC-ROC versus feature dimensionality. NucXplore achieves a mean AUC-ROC of 0.885 using 129 features.

**Supplementary Figure 7: Unsupervised structure and supervised performance in the LC25000 lung-and-colon cohort**

**A,** Median within-cluster inertia across repeated analyses for k = 2-8; shading shows the 95% interval. The modal elbow occurs at k = 4, compared with five annotated classes. **B,** Principal-component projection of the broader two-cluster solution. PC1 and PC2 explain 19.0% and 16.4% of the variance, respectively; black crosses denote cluster centres. **C,** Fold-wise accuracy, weighted precision, weighted recall, weighted F1 score, weighted one-versus-rest AUC-ROC, Cohen’s κ and MCC for the seven representations under five-fold stratified cross-validation. **D,** Macro-average and class-specific one-versus-rest ROC curves. NucXplore achieves AUC-ROC of 0.953 for the macro average, 0.918 for colon adenocarcinoma, 0.962 for lung adenocarcinoma, 0.991 for normal lung, 0.969 for lung squamous-cell carcinoma and 0.928 for normal colon.

**Supplementary Figure 8: Training and evaluation of skin cell-type and age-condition classifiers**

**A,** Distribution of n = 4,479 annotated nuclei across eight skin cellular compartments from 15 individuals and 123 image tiles. **B,** Three-dimensional PCA of the annotated nuclei, colored by cell type. PC1, PC2 and PC3 explain 18.6%, 15.2% and 13.6% of the variance, respectively. **C,** Training and held-out test performance of logistic regression, decision tree, random forest, extra trees, support vector machine, k-nearest neighbors, naive Bayes, multilayer perceptron, XGBoost and gradient boosting classifiers across seven evaluation metrics. **D,** Training and held-out test performance of the selected XGBoost cell-type classifier. **E,** Five-fold cross-validation distributions for accuracy, precision, recall, F1 score, Cohen’s κ, MCC and AUC-ROC for the selected XGBoost cell-type classifier. **F,** Row-normalised held-out-test confusion matrix for the selected eight-class XGBoost cell-type classifier. Rows and columns denote true and predicted labels, respectively; values show the percentage of each true class assigned to each predicted class. **G,** Mean absolute Shapley additive explanation (SHAP) values for representative cell-identity features across cell types. **H,** Transformer-based multiple-instance learning (MIL) architecture. Mini-bags of nucleus features are processed by a nucleus encoder and transformer, followed by gated-attention bag-level and instance-level heads; bag and instance focal losses are combined during optimization. **I,** Nucleus counts across six age brackets and eight cell types in SE and NSE skin. Inner rings show age-bracket totals and outer rings show the corresponding cell-type composition. SE, sun-exposed; NSE, non-sun-exposed. **J,** ROC curves for the six-class age model and two-class condition model. Age-class AUC-ROC range from 0.900 to 1.000; condition AUC-ROC are 1.000 for SE and 0.948 for NSE. **K,** Overall performance of the combined 12-class classifier. The panel reports accuracy, 0.7682; balanced accuracy, 0.8034; F1 macro, 0.7771; F1 weighted, 0.7729; precision macro, 0.7851; recall macro, 0.8034; AUC macro, 0.9818; MCC, 0.7446; and Cohen’s κ, 0.7393. **L,** Normalised importance of age-associated NucXplore features in independently trained SE and NSE models. **M,** Signed differential importance of features preferentially weighted by the SE or NSE age models. Positive values denote photoaging-enriched features and negative values denote intrinsic-aging-enriched features. These differences describe condition-associated model weighting and are not causal exposure effects. **N,** Mean shift in predicted age for cross-condition prediction. NSE-to-SE denotes predictions for NSE samples using the SE model, whereas SE-to-NSE denotes predictions for SE samples using the NSE model. color indicates the shift in years for each true age bracket. **O,** Principal-component projections of age-probability profiles, colored by chronological age bracket or exposure condition.

**Supplementary Figure 9: Cell-type-resolved performance of combined age-condition classification**

**A,** One-versus-rest ROC curves for the 12 combined age-condition classes within each of the eight skin cell types. SE and NSE states are shown for each age bracket from 20-29 to 70-79 years; class-specific AUC-ROC are listed within each plot. SE, sun-exposed; NSE, non-sun-exposed. **B,** Row-normalised confusion matrix for the combined 12-class model. Rows and columns denote true and predicted age-condition states, respectively, and values show the fraction of each true class assigned to each predicted class.

**Supplementary Figure 10: Sample-preserving subsampling, model training and condition-dependent nuclear** aging dynamics

**A,** Two-level, sample-preserving workflow based on FAISS (Facebook AI Similarity Search) used to select a representative subset of the full GTEx nucleus-feature dataset. Data are stratified by cell type, age bracket and condition; local per-sample selection preserves sample representation, followed by global refinement to maintain feature diversity and prevent sample dropout. **B,** Age-bracket and cell-type composition of the selected SE and NSE subsets. Inner rings show age-bracket totals and outer rings show cell-type composition. SE, sun-exposed; NSE, non-sun-exposed. **C,** Coverage of all age-bracket-by-cell-type strata in the selected subsets for both conditions. **D,** Feature-wise Kolmogorov-Smirnov (KS) statistics comparing the selected subsets with their corresponding source distributions. **E,** Raw and smoothed flow-matching training loss over 100 epochs for each cell-type model. **F,** Mean feature-space speed across model age for each cell type under SE and NSE conditions. Bars show the age-grid mean and error bars show the descriptive SEM across the 51 model-age positions; these are not donor-level uncertainty estimates. **G,** Distributions of feature-level standardized effects from the condition-switch simulation for each cell type. **H,** RMS feature-velocity trajectories for keratinocytes, hair follicles, sebaceous glands, sweat glands, arrector pili and blood vessels. Solid and dashed lines denote SE and NSE conditions. Panels report the mean SE-minus-NSE difference and a heteroscedasticity- and autocorrelation-consistent comparison with Holm-adjusted P value; none of the six comparisons is significant after correction.

**Supplementary Figure 11: Source-free adaptation and region-level predictions in the external hospital cohort**

**A,** Per-sample, per-cell-type adaptation workflow. Viable nuclei are assembled into mini-bags, followed by source hypothesis transfer (SHOT), a source-free unsupervised adaptation method, for the age and condition models; two-stage inference, validation-F1-weighted ensembling and export of adapted models and predictions then follow. **B,** Region-level predictions for 24 regions of interest from eight individuals. Columns report chronological age, sex, true and predicted age bracket, marginal expected age, expected NSE age, NSE age error, NSE probability and nucleus count. The cohort contains 2,399,803 nuclei and the region-level MAE is 5.9 years. NSE, non-sun-exposed. **C,** Expected NSE age for individual regions of interest, grouped by patient. Symbols denote chronological age, predicted SE and NSE ages, and the nucleus-weighted patient-level expected NSE age **D,** Cell-type-specific condition-classification accuracy, ranging from 17% for sweat-gland nuclei to 96% for hair-follicle nuclei and 100% for keratinocytes.

## SUPPLEMENTARY DATA

**Supplementary Table 1: NucXplore feature definitions**

Complete list of the 129 nuclear phenomic features implemented in NucXplore, organized across morphology, chromatin, intensity, texture, color, and spatial-position domains, together with the corresponding feature names and definitions.

**Supplementary Table 2: Software packages and computational environment**

Software, libraries, and package versions used for NucXplore feature extraction, image processing, machine-learning analyses, statistical evaluation, and downstream computational workflows.

**Supplementary Table 3: Benchmarking performance in the Borkowski *et al.* dataset**

Performance of NucXplore and the comparator representations in the lung and colon histopathology dataset from Borkowski *et al.*, including classification metrics used to evaluate discrimination across the five histological classes.

**Supplementary Table 4: Benchmarking performance in the Freire *et al.* dataset**

Performance of NucXplore and the comparator representations in the oral squamous cell carcinoma dataset from Freire *et al.*, including classification metrics for distinguishing cancer-associated and non-cancer nuclear populations.

**Supplementary Table 5: Benchmarking performance in the Schuiveling *et al.* dataset**

Performance of NucXplore and the comparator representations in the melanoma dataset from Schuiveling *et al.*, including multiclass classification metrics across the annotated nuclear phenotypes.

**Supplementary Table 6: Skin cell-type classification performance**

Classification performance of the NucXplore-based skin-compartment model across eight major cellular populations: adipocytes, arrector pili, blood vessels, fibroblasts, hair follicles, keratinocytes, sebaceous glands, and sweat glands.

**Supplementary Table 7: Age and exposure classification performance for adipocytes**

Performance of the adipocyte-specific hierarchical models for chronological age, sun-exposure condition, and joint age-exposure classification, evaluated using complementary classification metrics.

**Supplementary Table 8: Age and exposure classification performance for arrector pili cells**

Performance of the arrector-pili-specific hierarchical models for chronological age, sun-exposure condition, and joint age-exposure classification.

**Supplementary Table 9: Age and exposure classification performance for blood-vessel nuclei**

Performance of the blood-vessel-specific hierarchical models for chronological age, sun-exposure condition, and joint age-exposure classification.

**Supplementary Table 10: Age and exposure classification performance for fibroblasts**

Performance of the fibroblast-specific hierarchical models for chronological age, sun-exposure condition, and joint age-exposure classification.

**Supplementary Table 11: Age and exposure classification performance for hair-follicle nuclei**

Performance of the hair-follicle-specific hierarchical models for chronological age, sun-exposure condition, and joint age-exposure classification.

**Supplementary Table 12: Age and exposure classification performance for keratinocytes**

Performance of the keratinocyte-specific hierarchical models for chronological age, sun-exposure condition, and joint age-exposure classification.

**Supplementary Table 13: Age and exposure classification performance for sebaceous-gland nuclei**

Performance of the sebaceous-gland-specific hierarchical models for chronological age, sun-exposure condition, and joint age-exposure classification.

**Supplementary Table 14: Age and exposure classification performance for sweat-gland nuclei**

Performance of the sweat-gland-specific hierarchical models for chronological age, sun-exposure condition, and joint age-exposure classification.

**Supplementary Table 15: Cross-condition age-prediction analysis**

Summary of age predictions obtained under matched and cross-condition evaluation between sun-exposed and non-sun-exposed skin, showing how exposure condition influences predicted morphological age across chronological age groups.

**Supplementary Table 16: Conditional flow-matching model training losses**

Summary of the training-loss statistics for the cell-type-specific conditional flow-matching models used to reconstruct continuous nuclear phenotypic trajectories across the adult age range.

**Supplementary Table 17: Conditional flow-matching model configuration**

Architecture, optimization parameters, training settings, and numerical integration configuration used for the cell-type-specific conditional flow-matching and Neural ODE models.

**Supplementary Table 18: Conditional flow-matching trajectory summary**

Summary statistics derived from the fitted cell-type-specific conditional flow-matching models, describing the modeled nuclear phenotypic trajectories across age and sun-exposure conditions.

**Supplementary Table 19: Cross-cell-type comparison of nuclear phenotypic dynamics**

Comparison of model-derived nuclear phenotypic dynamics across the eight skin cell types and exposure conditions, summarizing differences in the magnitude of morphological change across the modeled lifespan.

**Supplementary Table 20: Age-bin statistics for nuclear phenotypic velocity and acceleration**

Age-resolved summary statistics for model-derived nuclear phenotypic velocity and acceleration in sun-exposed and non-sun-exposed skin, used to characterize changes in the rate of nuclear morphological aging across the adult lifespan.

## Notes

### Competing Interest Statement

The authors have declared no competing interest.

