## Supplementary Figure 1 for "Scalable Computational Phenomics of Nuclear Morphology"

## A

| Domain | Features (code names) | Mathematical definition | Biological significance |
|---|---|---|---|
| Morphological - Size | area, perimeter, equivalent_diameter, convex_area | $A = \Sigma\ 1$ over $p \in M$<br>$P = \Sigma\ \lVert x_{i+1} - x_i \rVert$ ($i$ = 1..N−1)<br>$d_{eq} = \sqrt{(4A/\pi)}$<br>$A_{conv}$ = area(ConvexHull($M$))<br>*where:* $M$ = binary nuclear mask; $p$ = pixel index; $A = \lvert M \rvert$; $x_i$ = ordered boundary point $i$. | Absolute nuclear size, boundary length, and reference convex enclosure. Reflect DNA content, ploidy, and overall cell-cycle / transcriptional state; define the geometric scale against which envelope deformations are measured. |
| Morphological - Shape (regularity) | eccentricity, solidity, extent, circularity, aspect_ratio | $e = \sqrt{(1 - b^2/a^2)}$<br>$S = A/A_{conv}$; $E_x = A/A_{bbox}$<br>$C = 4\pi A/P^2$; $AR = a/b$<br>*where:* $a \geq b$ = semi-axes of the ellipse whose 2nd central moments match $M$; $A_{bbox}$ = axis-aligned bounding-box area. | Boundary regularity, compactness, and elongation of the nucleus. Classical histopathological descriptors of nuclear shape that report cytoskeletal forces, lamin-scaffold integrity, and cell polarity within the surrounding tissue. |
| Morphological - Shape (axes / topology) | major_axis_length, minor_axis_length, orientation, euler_number | $L_a = 2a$; $L_b = 2b$<br>$\theta \in [-\pi/2, \pi/2]$ = angle of $a$ w.r.t. horizontal<br>$\chi = N_c - N_h$<br>*where:* $N_c$ = # connected components in $M$; $N_h$ = # holes in $M$. | Directional extents, tissue alignment, and topological integrity. Quantify whether the nucleus is isotropically scaled or directionally deformed; $\chi < 1$ indicates intranuclear vacuoles or accompanying micronuclei. |
| Morphological - Hu moments | hu_moment_1, hu_moment_2, hu_moment_3, hu_moment_4, hu_moment_5, hu_moment_6, hu_moment_7 | $\mu_{pq} = \Sigma\Sigma\ (x - x)^p\ (y - \bar{y})^q\ I(x,y)$<br>$\eta_{pq} = \mu_{pq}\ /\ \mu_0\ ^{(p+q)/2 + 1}$<br>$\varphi_1 = \eta_{20} + \eta_{02}$<br>$\varphi_2 = (\eta_{20} - \eta_{02})^2 + 4\eta_{11}{}^2$<br>$\varphi_3 \ldots \varphi_7$ : standard Hu polynomials in $\eta_{pq}$<br>*where:* $(x, \bar{y})$ = intensity-weighted centroid; $I(x,y)$ = pixel intensity; $\lvert\varphi_k\rvert$ capped at $10^{10}$; $\varphi_7$ uniquely changes sign under mirror reflection. | Seven translation-, rotation- and scale-invariant shape descriptors. Provide geometry-invariant shape characterisation usable across cell types, magnifications and tissue orientations; $\varphi_7$ isolates chirality of the nuclear contour. |
| Morphological - Advanced shape | convexity, fractal_dimension, roughness, bending_energy | $C_x = A_{conv}/A$; $D_f = \log P / \log A$<br>$R = P^2/A$; $BE = \Sigma\ \kappa_i{}^2$<br>$\kappa = (x'y'' - y'x'')\ /\ (x'^2 + y'^2)^{3/2}$<br>*where:* $(x(t), y(t))$ = contour parametrisation (skimage.find_contours @ 0.5); primes = derivatives w.r.t. $t$; $\lvert\kappa\rvert$ capped at $10^6$. | Contour tortuosity, self-similar boundary complexity, and stored envelope bending energy. Together describe how far the envelope departs from a smooth ellipse — through indentations, multi-scale complexity, and integrated micro-curvature. |
| Morphological - Fourier | fourier_descriptor_1, fourier_descriptor_2, fourier_descriptor_3, fourier_descriptor_4, fourier_descriptor_5 | $z(t) = x(t) + i\ y(t)$<br>$Z_k = \mathrm{FFT}[z(t)]$<br>$FD_k = \lvert Z_k \rvert\ /\ \lvert Z_0 \rvert$, $k = 1 \ldots 5$<br>*where:* $z(t)$ = complex contour signal; extraction via cv2.findContours (CHAIN_APPROX_NONE); normalisation by $\lvert Z_0 \rvert$ makes $FD_k$ scale-invariant. | Low- to mid-frequency boundary undulations of the nuclear contour, indexed by harmonic mode. $FD_1$–$FD_2$ encode dominant elliptical and elongation modes; $FD_3$–$FD_5$ resolve progressively finer multi-lobed and crenulated boundary detail. |
| Morphological - NEIS | neis_irregularity_score, neis_spectral_energy, neis_spectral_peak_mode | $r(\theta)$ = centroid-to-contour radius at 64 uniform angles<br>$F_k = \mathrm{FFT}[r(\theta)]$<br>$\mathrm{NEIS} = (\Sigma\ \lvert F_k \rvert^2$ for $k \geq 6)\ /\ (\Sigma\ \lvert F_k \rvert^2$ for $k$ = 2..5)<br>$SE = \Sigma\ \lvert F_k \rvert^2$ for $k \geq 1$; $k^* = \mathrm{argmax}\ \lvert F_k \rvert^2$ for $k \geq 4$ | Compact spectral fingerprint of nuclear-envelope irregularity: high-to-low frequency ratio, total non-DC power, and dominant high-frequency mode. Distinguishes smooth contours from those with characteristic micro-undulations of the lamina; the peak mode identifies the spatial scale of undulation. |
| Positional | centroid_row, centroid_col, distance_to_nearest_neighbor | $c_r = (1/A)\ \Sigma\ r_p$; $c_c = (1/A)\ \Sigma\ c_p$ (full-WSI coords)<br>$NND_i = \min \lVert c_i - c_j \rVert$ for $j \neq i$<br>*where:* $r_p$, $c_p$ = row / column of pixel $p$; $c_i$ = centroid of nucleus $i$; NND via scipy.spatial.distance.cdist. | Tissue-level position of each nucleus and distance to its closest neighbor. Anchor nuclei in tissue coordinates and report local cell-packing density; support analyses of epithelial stratification and overall tissue cellularity. |
| Intensity - 1st / 2nd order | pre_norm_/post_norm_ × {mean_intensity, median_intensity, std_intensity, min_intensity, max_intensity, range_intensity, iqr_intensity} | $\mu = (1/N)\ \Sigma\ I_p$; $\tilde{I} = Q_2$; $\sigma = \sqrt{[(1/N)\ \Sigma\ (I_p - \mu)^2]}$<br>$I_{min} = \min I_p$; $I_{max} = \max I_p$<br>$R_I = I_{max} - I_{min}$; $IQR = Q_3 - Q_1$<br>*where:* $I_p$ = grayscale intensity at pixel $p$; $N = \lvert M \rvert$; $Q_1$, $Q_2$, $Q_3$ = distribution quartiles. | Location and spread of the intra-nuclear intensity distribution. Report bulk chromatin compaction (mean / median), the dynamic range of compaction states within a nucleus (range / IQR), and identify extreme dense or relaxed points. |
| Intensity - Higher moments | pre_norm_/post_norm_ × {skewness_intensity, kurtosis_intensity, entropy_intensity} | $\gamma_1 = \mathbb{E}\quad I - \mu)/\sigma)^3]$<br>$\gamma_2 = \mathbb{E}\quad I - \mu)/\sigma)^4] - 3$<br>$H = -\ \Sigma\ p_i \log_2 p_i$ over 256-bin histogram<br>*where:* $\mathbb{E}$ $M$; $p_i$ = normalised bin frequency; $\gamma_2$ is excess kurtosis (0 for Gaussian). | Asymmetry, tail weight, and information-theoretic disorder of the intensity histogram. Reveal whether the chromatin distribution is balanced, skewed toward dense or relaxed states, or contains rare extreme foci such as condensed patches or repair bodies. |
