## Supplementary Figure 2 for "Scalable Computational Phenomics of Nuclear Morphology"

**A**

| Domain | Features (code names) | Mathematical definition | Biological significance |
|---|---|---|---|
| Texture - GLCM | pre_norm_/post_norm_ × {glcm_contrast, glcm_dissimilarity, glcm_homogeneity, glcm_energy, glcm_correlation, glcm_ASM} | $P(i,j)$ = normalised GLCM, $d$ = 1, θ ∈ {0°, 45°, 90°, 135°} (averaged); 256 grey levels<br>Contrast = $\Sigma (i - j)^2 P$; Dissim = $\Sigma \lvert i - j \rvert P$<br>Homog = $\Sigma P/(1 + \lvert i - j \rvert)$; Energy = $\sqrt{(\Sigma P^2)}$;<br>ASM = $\Sigma P^2$<br>Corr = $\Sigma (i - \mu_i)(j - \mu_j) P / (\sigma_i \sigma_j)$<br>*where:* $i, j$ = paired intensity values; $\mu_i, \sigma_i$ = row-marginal moments of $P$. | Local co-occurrence statistics of paired intensity values — transition magnitude, uniformity, and directional dependence. Quantify fine-scale chromatin organisation: magnitude of local density transitions, uniformity of texture, and directionality of chromatin patterns. |
| Texture - LBP | pre_norm_/post_norm_ × {lbp_mean, lbp_std, lbp_entropy} | LBP($x_c$) = $\Sigma s(I_p - I_c) \cdot 2^p$ ($p$ = 0..7)<br>$s(u)$ = 1 if $u \geq 0$ else 0; P = 8, R = 1, uniform → 10 bins<br>mean L, std $\sigma_L$ over $M$; $H_L = - \Sigma p_b \log_2 p_b$ ($b$ = 1..10) | Rotationally tolerant micro-texture of chromatin patches — dominant local pattern, diversity, and disorder. Characterise which kinds of local chromatin micro-pattern (flat, edge, corner) dominate the nucleus, and how varied or disordered those patterns are. |
| Texture - HOG | pre_norm_/post_norm_ × {hog_mean, hog_std, hog_max, hog_min} | $v$ = HOG($I \cdot \mathbb{1}$): $\nabla I$ → 8 orientation bins per 8×8-px cell, L2-normalised per cell<br>v, $\sigma_v$, $v_{max}$, $v_{min}$ = mean / std / max / min of the concatenated HOG vector<br>*where:* $\mathbb{1}$ = mask indicator function. | Strength and heterogeneity of internal chromatin edges. Report the sharpness and distribution of intra-nuclear chromatin boundaries — distinguishing smoothly textured nuclei from those with focal high-gradient edges. |
| Color - Hematoxylin | pre_norm_/post_norm_ × {mean_hematoxylin, std_hematoxylin, skew_hematoxylin, kurt_hematoxylin, min_hematoxylin, max_hematoxylin} | OD = − log(($I$ + ε)/255); [$H_p$, $E_p$] = $\mathbf{S}^{-1}$ OD$_p$<br>$\mu_H, \sigma_H, \gamma_{1,H}, \gamma_{2,H}, H_{min}, H_{max}$ : same formulas as Intensity block, applied to $H_p$<br>*where:* **S** = Ruifrok–Johnston stain matrix; H-vec = [0.65, 0.70, 0.29]; E-vec = [0.07, 0.99, 0.11]; $H_p$, $E_p$ = per-pixel hematoxylin / eosin concentration. | Distribution of stain-deconvolved chromatin-binding signal. Direct readout of DNA / nucleic-acid density isolated from cytoplasmic protein staining — the preferred chromatin-specific intensity channel. |
| Color - Eosin | pre_norm_/post_norm_ × {mean_eosin, std_eosin, skew_eosin, kurt_eosin, min_eosin, max_eosin} | $\mu_E, \sigma_E, \gamma_{1,E}, \gamma_{2,E}, E_{min}, E_{max}$ : same formulas as Intensity block, applied to $E_p$ | Distribution of residual cytoplasmic / protein signal inside the nucleus. Should be near zero in intact nuclei; elevated intra-nuclear eosin indicates protein inclusions, envelope leakage, or partial-volume contamination by adjacent cytoplasm. |
| Color - H:E ratio | pre_norm_/post_norm_ × {he_ratio_H_to_E} | $R_{H/E} = \mu_H / \mu_E$ (safe division: returns 0 if $\mu_E$ = 0) | Chromatin-to-protein staining balance per nucleus. Compact single-number indicator of nuclear composition — high values reflect chromatin-dominated nuclei, low values reflect protein-rich or eosin-contaminated nuclei. |
| Chromatin - CCSM (area / count) | pre_norm_/post_norm_ × {ccsm_condensed_area_ratio, ccsm_num_clumps} | $\tilde{I}$ = CLAHE($I$; clip = 0.03, kernel = 16)<br>$M_{cond}$ ← argmin-cluster of GMM$_{k=2}$($\tilde{I}$) (darker component)<br>then opening + remove-small (min_size = 10)<br>$\rho = A_{cond} / A_{nuc}$; $K$ = $\lvert$connected components of $M_{cond}\rvert$ | Fraction of the nucleus that is condensed and count of discrete condensed foci. Direct quantification of heterochromatin content; distinguish nuclei with few large heterochromatin domains from those with many small fragmented foci. |
| Chromatin - CCSM (clump morphology) | pre_norm_/post_norm_ × {ccsm_mean_clump_area, ccsm_mean_clump_eccentricity, ccsm_mean_clump_solidity} | $\bar{A}_c = (1/K) \Sigma A_k$; $\bar{e}_c = (1/K) \Sigma e_k$; $S_c = (1/K) \Sigma S_k$<br>*where:* $A_k, e_k, S_k$ = area / eccentricity / solidity of clump $k$ via regionprops. | Mean size, elongation, and regularity of individual heterochromatin domains. Describe whether condensation tracks compact isotropic blobs, elongated (e.g., lamina-associated) structures, or irregular fragmented configurations. |
| Chromatin - CCSM (clump topology) | pre_norm_/post_norm_ × {ccsm_mean_dist_to_boundary, ccsm_mean_nnd} | $d_b = (1/\lvert M_{cond}\rvert) \Sigma EDT(M)\rvert_p$ for $p \in M_{cond}$<br>$d_{nn} = (1/K) \Sigma \min \lVert c_i - c_j \rVert$ ($j \neq i$)<br>*where:* EDT via scipy.ndimage.distance_transform_edt; $c_i$ = centroid of clump $i$. | Radial position and inter-cluster spacing of heterochromatin within the nucleus. Discriminate peripheral, lamina-associated heterochromatin from interior heterochromatin, and report whether condensed domains are evenly dispersed or clustered. |
| Chromatin - CCSM (intra-clump texture) | pre_norm_/post_norm_ × {ccsm_contrast, ccsm_correlation, ccsm_energy, ccsm_homogeneity} | $P_{ccsm}$ = GLCM at $d$ = 1, θ = 0 on $\tilde{I} \cdot \mathbb{1}$<br>Contrast, Correlation, Energy, Homogeneity computed as in Texture — GLCM but restricted to the condensed region. | Local co-occurrence texture restricted to the heterochromatic compartment. Characterise the internal sub-structure of heterochromatin — directional organisation and textural uniformity — independently from the texture of the nucleus as a whole. |
