## Supplementary figures and images for "Scalable Computational Phenomics of Nuclear Morphology"

### Supplementary Figure 3

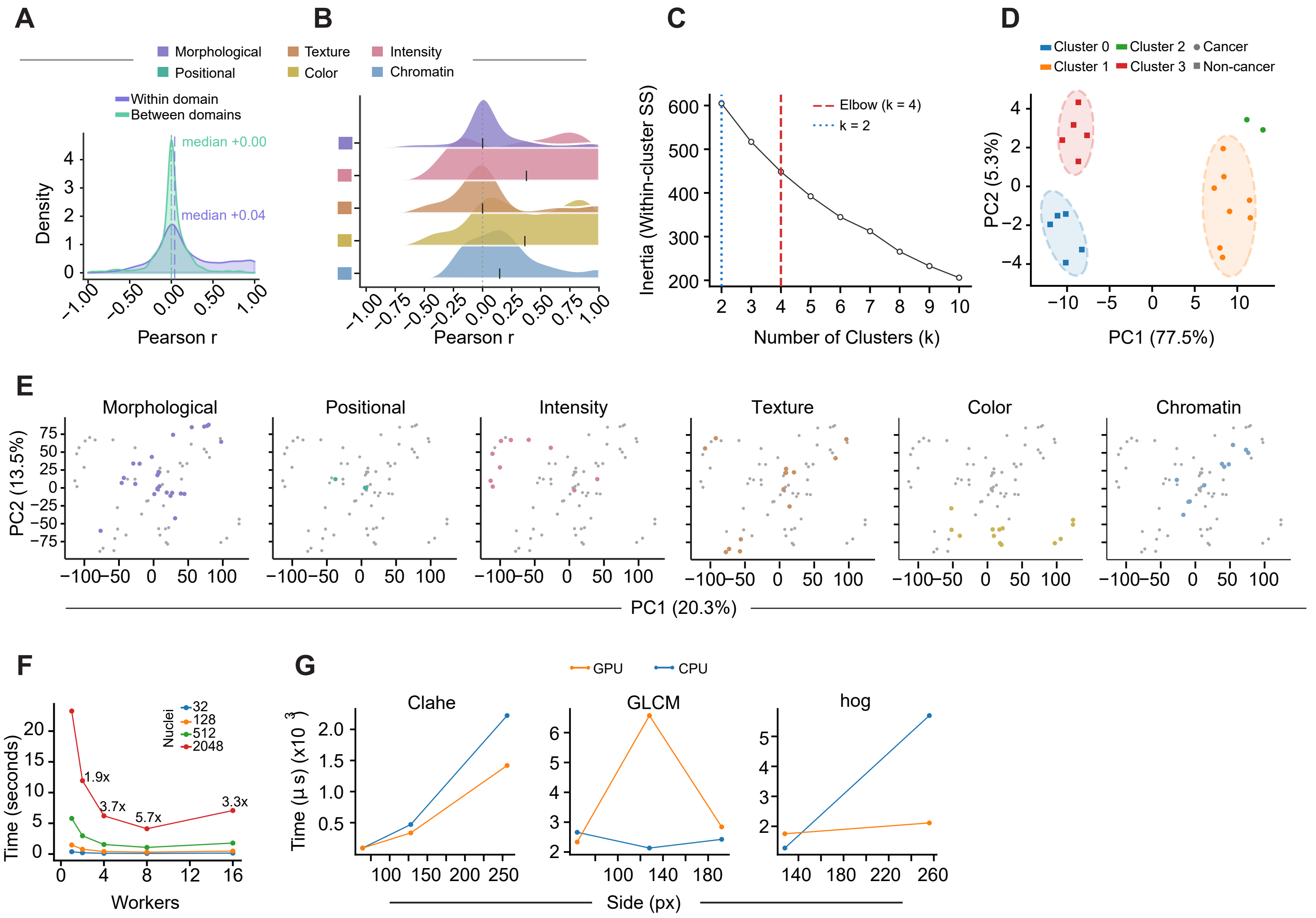

### Supplementary Figure 4

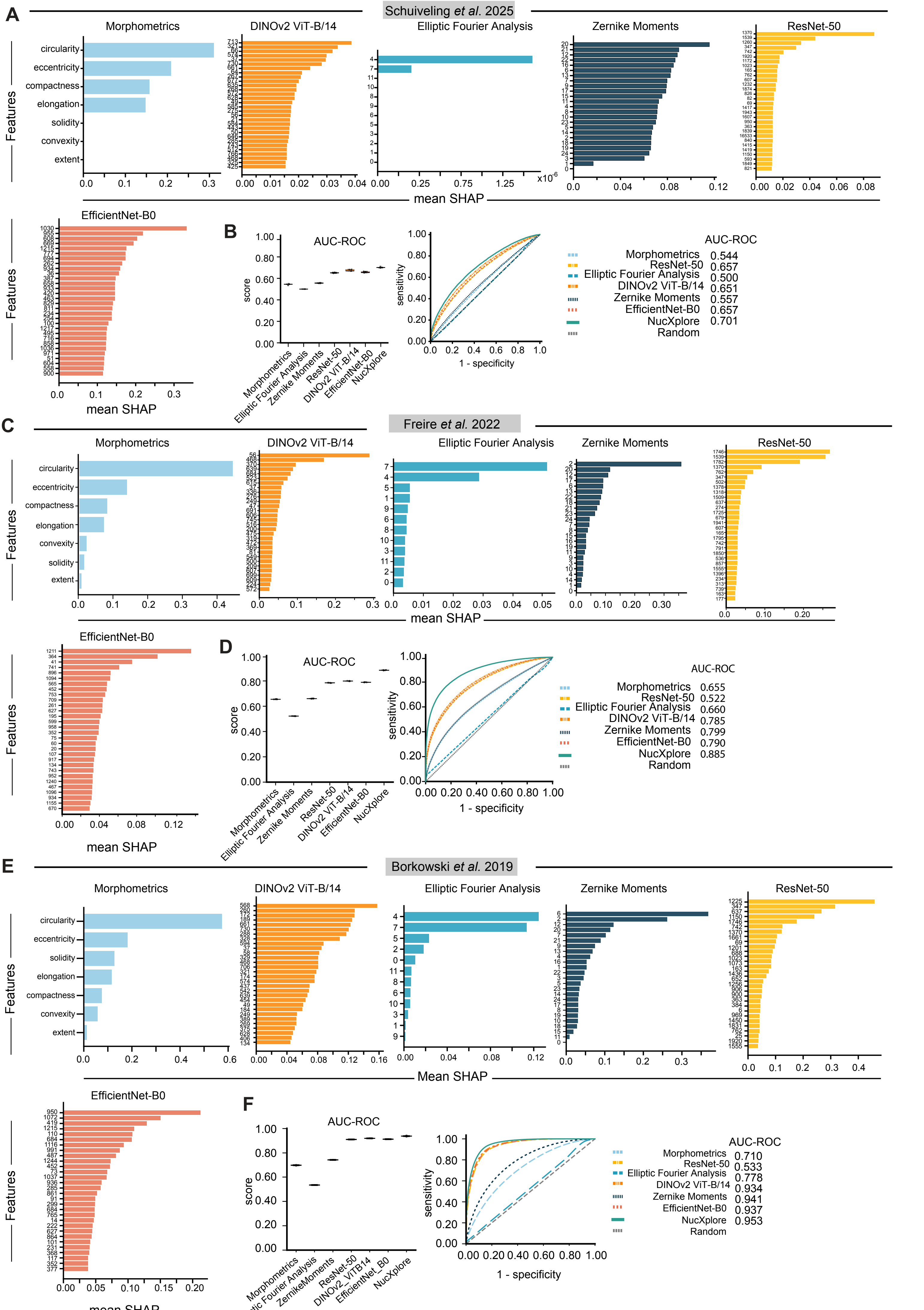


Supplementary Figure 4

### Supplementary Figure 5

Schuiveling *et al.* 2025

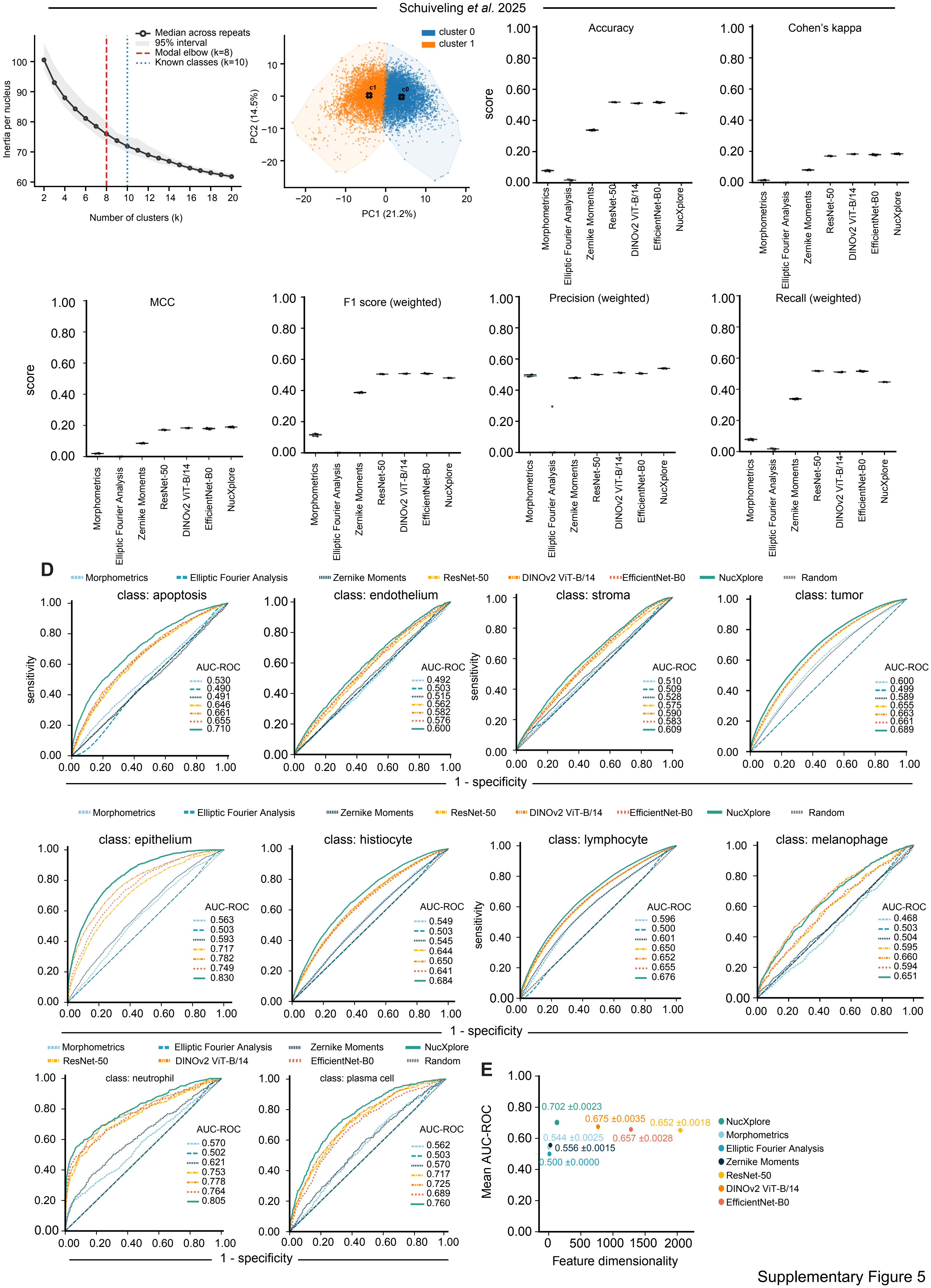


Supplementary Figure 5

### Supplementary Figure 6

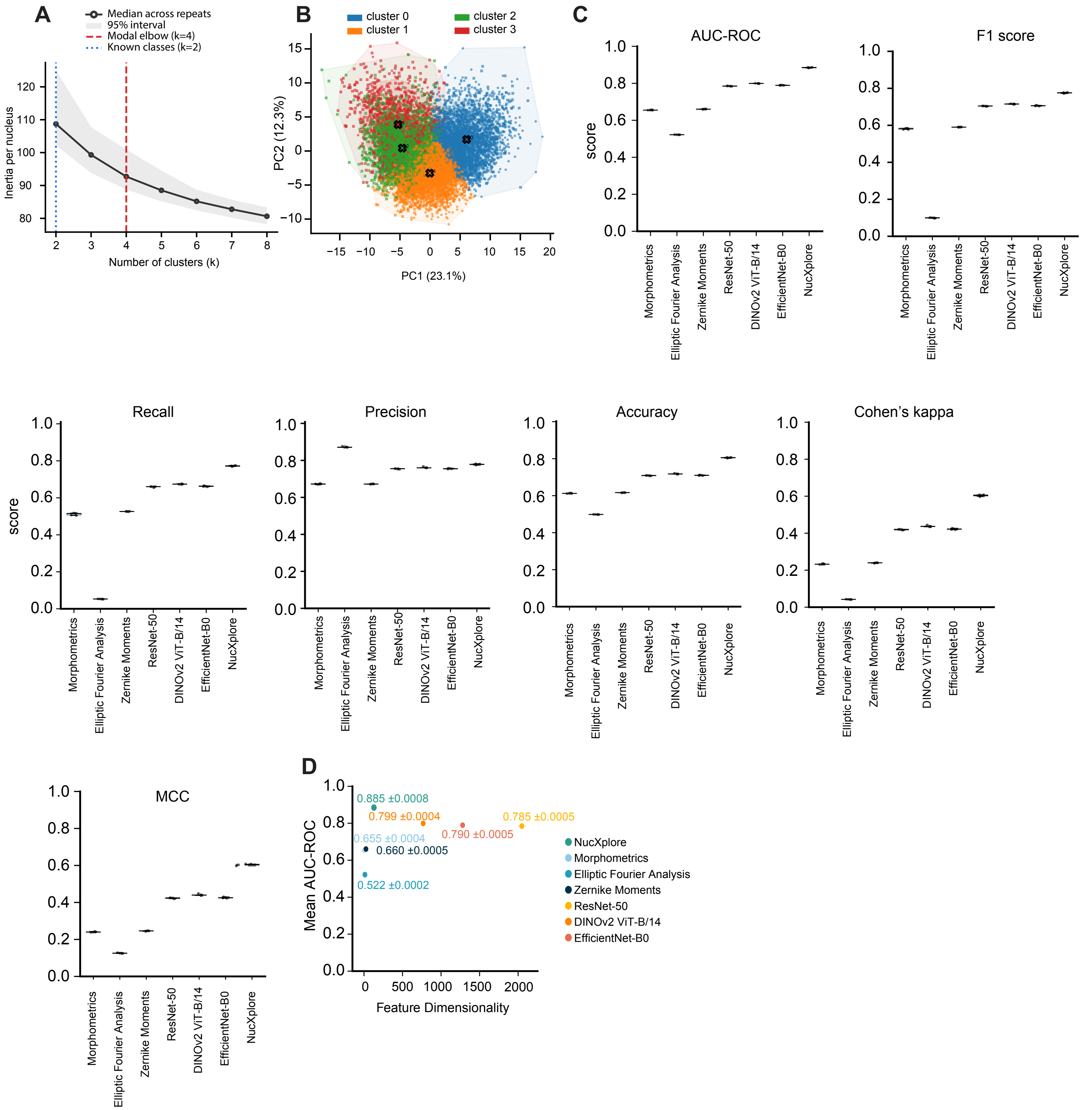

### Supplementary Figure 7

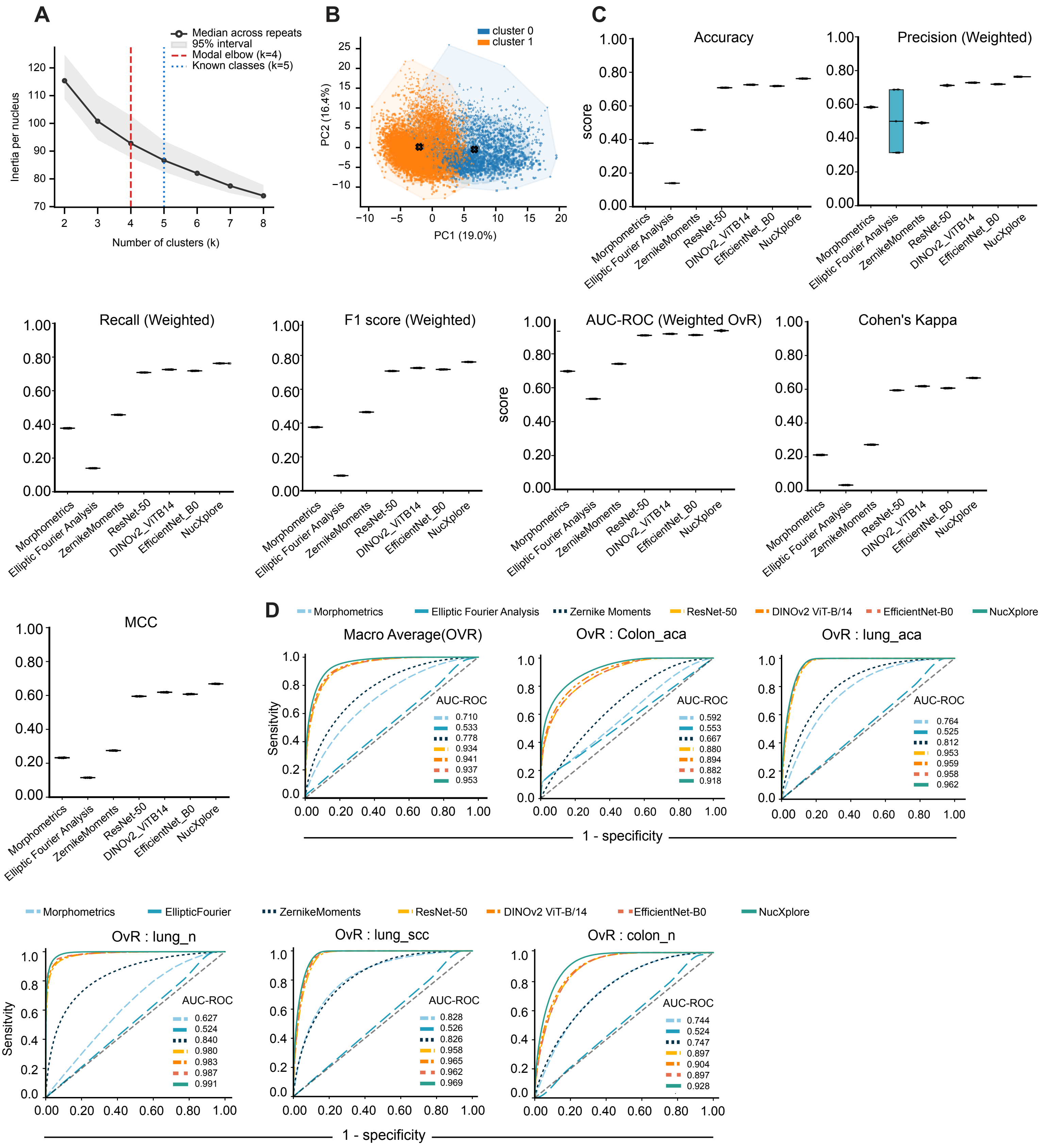


Supplementary Figure 7

### Supplementary Figure 8

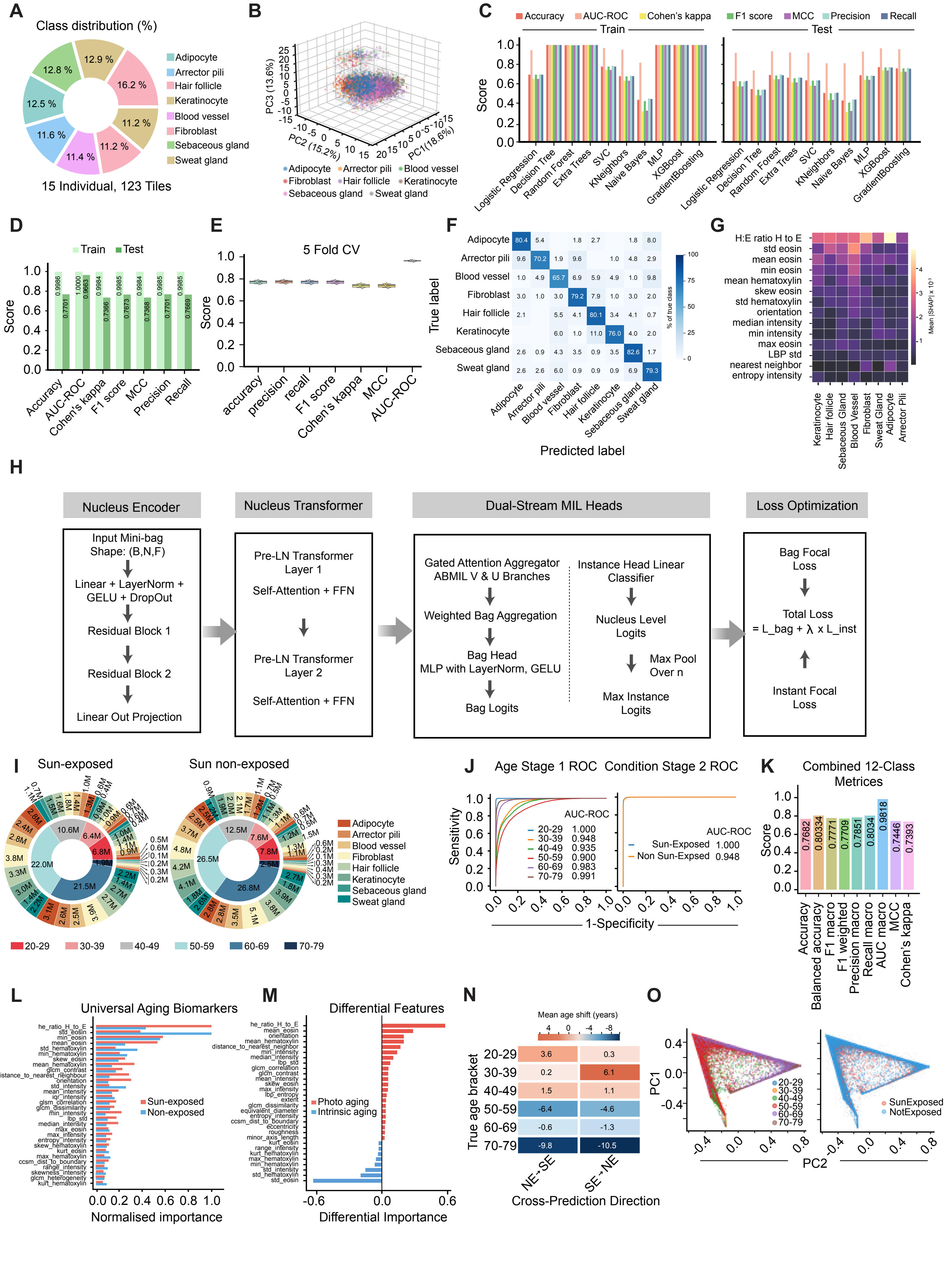


Supplementary Figure 8

### Supplementary Figure 9

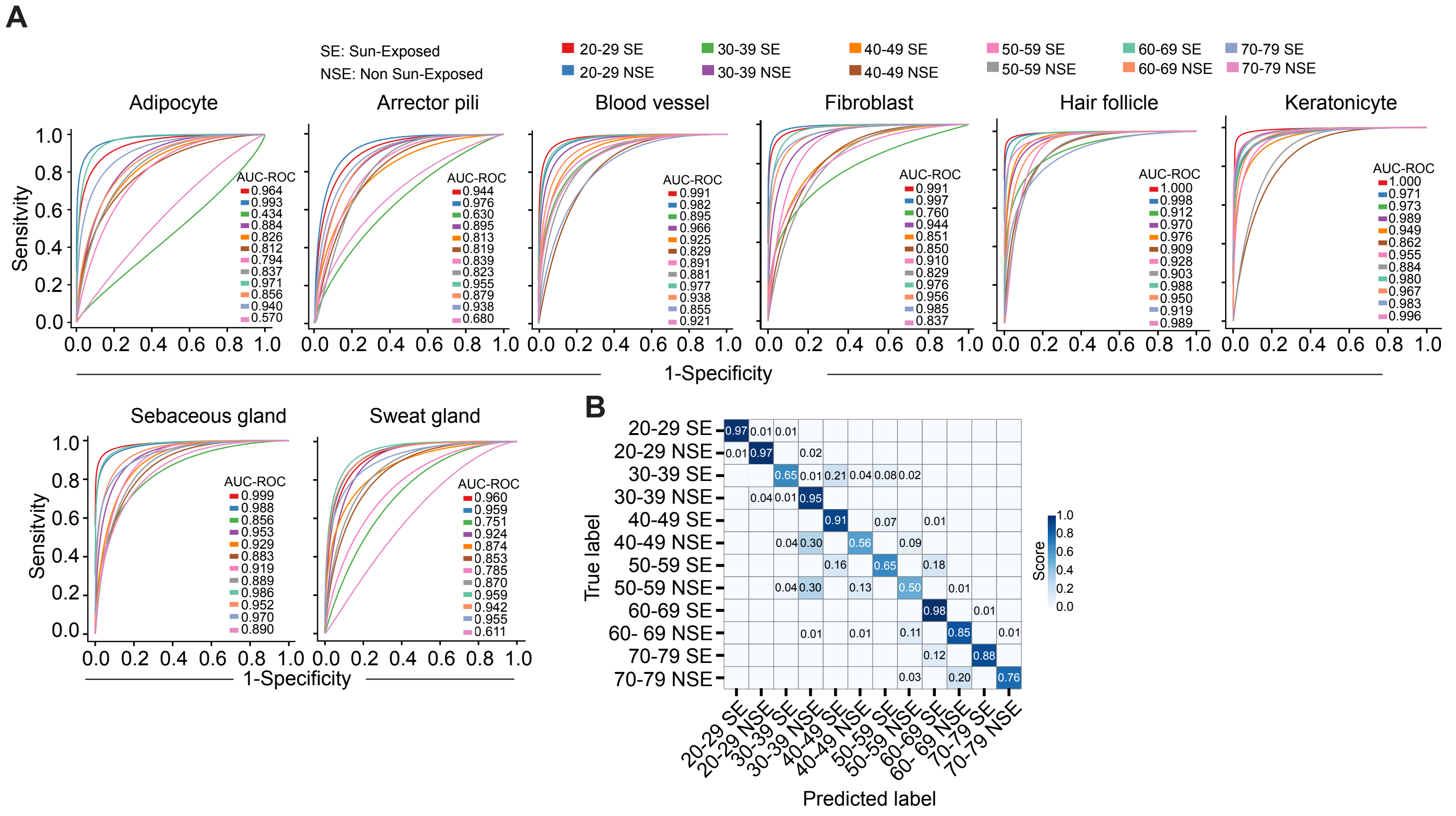


Supplementary Figure 9

### Supplementary Figure 10

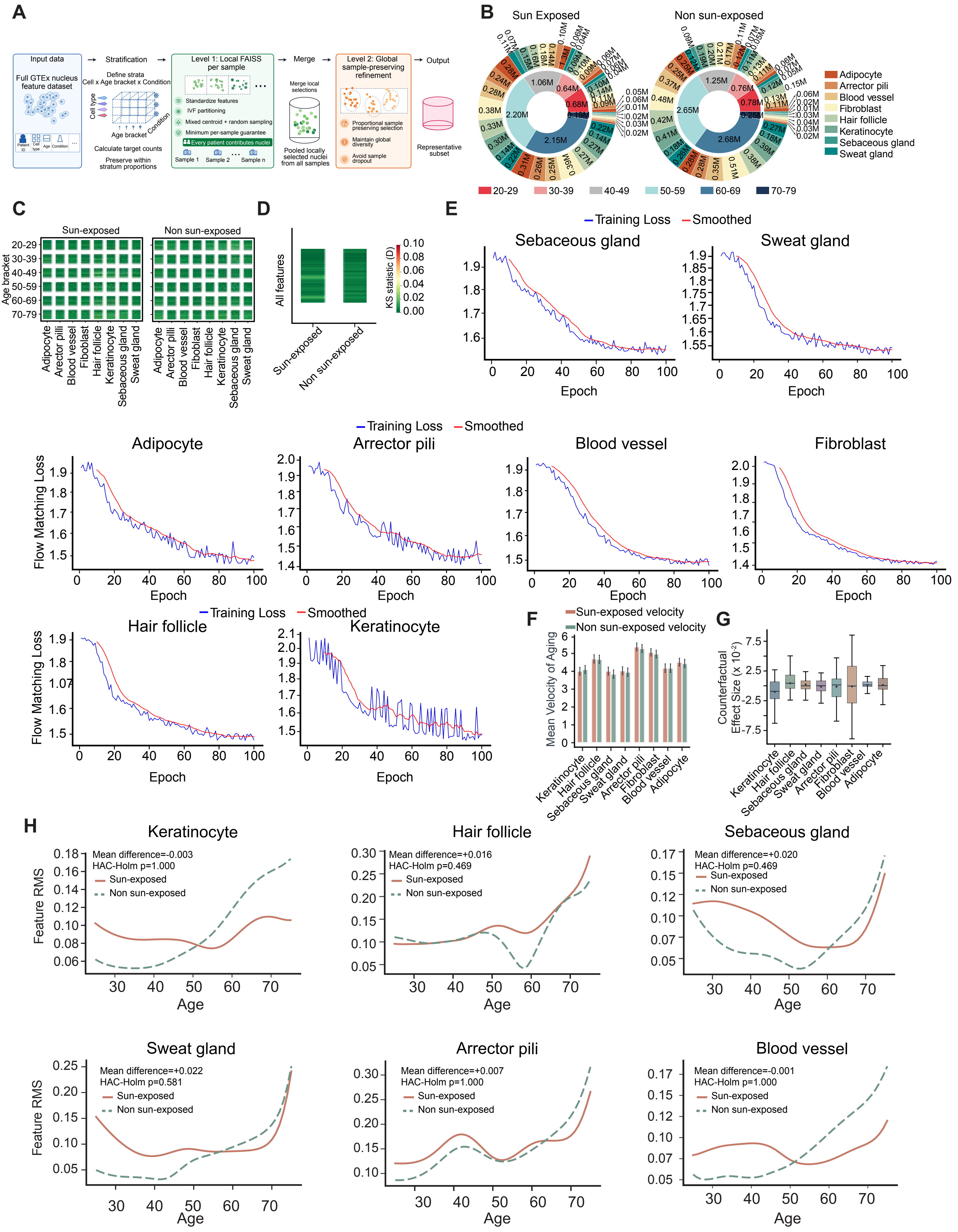


Supplementary Figure 10
