## Supplementary Figure 11 for "Scalable Computational Phenomics of Nuclear Morphology"

**A**

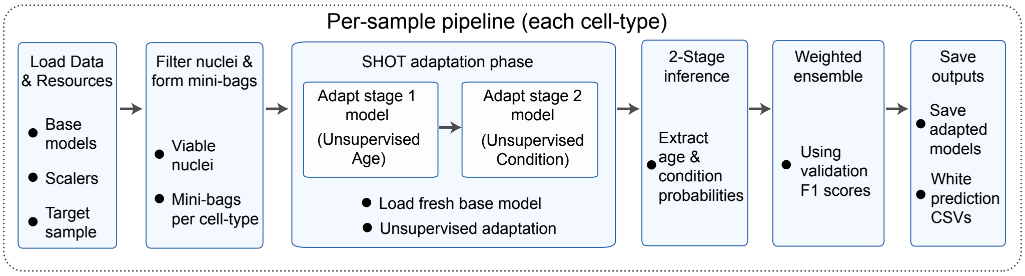


**B**

| Sample | True Age | Sex | True Bracket | Predicted | E[age] (marg) | E[age] (NE) | Err (NE) | P(NE) | Nuclei |
|---|---|---|---|---|---|---|---|---|---|
| C30-IMG1 | 34 | M | 30-39 | 30-39 | 48.9 | 44.0 | +10.0 | 0.560 | 71,973 |
| C30-IMG3 | 34 | M | 30-39 | 30-39 | 49.9 | 46.1 | +12.1 | 0.627 | 71,802 |
| C30-IMG5 | 34 | M | 30-39 | 50-59 | 46.7 | 41.0 | +7.0 | 0.336 | 70,849 |
| C29-IMG3 | 39 | F | 30-39 | 20-29 | 48.2 | 45.6 | +6.6 | 0.786 | 96,800 |
| C29-IMG4 | 39 | F | 30-39 | 30-39 | 47.7 | 39.6 | +0.6 | 0.679 | 89,575 |
| C29-IMG5 | 39 | F | 30-39 | 20-29 | 44.7 | 39.8 | +0.8 | 0.776 | 85,872 |
| C32-IMG1 | 43 | M | 40-49 | 30-39 | 48.9 | 47.0 | +4.0 | 0.777 | 42,342 |
| C32-IMG2 | 43 | M | 40-49 | 20-29 | 47.4 | 46.4 | +3.4 | 0.676 | 39,939 |
| C32-IMG3 | 43 | M | 40-49 | 50-59 | 55.0 | 54.7 | +11.7 | 0.684 | 31,875 |
| C31-IMG2 | 49 | M | 40-49 | 20-29 | 44.3 | 43.3 | -5.7 | 0.928 | 313,445 |
| C31-IMG3 | 49 | M | 40-49 | 30-39 | 50.9 | 49.4 | +0.4 | 0.907 | 300,382 |
| C31-IMG4 | 49 | M | 40-49 | 20-29 | 45.9 | 43.8 | -5.2 | 0.755 | 328,956 |
| C33-IMG2 | 49 | M | 40-49 | 60-69 | 62.1 | 61.2 | +12.2 | 0.794 | 2,795 |
| C33-IMG3 | 49 | M | 40-49 | 20-29 | 46.9 | 45.6 | -3.4 | 0.984 | 177,735 |
| C33-IMG5 | 49 | M | 40-49 | 70-79 | 50.9 | 50.2 | +1.2 | 0.928 | 178,395 |
| C34-IMG2 | 51 | M | 50-59 | 70-79 | 53.9 | 53.5 | +2.5 | 0.681 | 62,066 |
| C34-IMG4 | 51 | M | 50-59 | 70-79 | 52.7 | 52.3 | +1.3 | 0.803 | 58,433 |
| C34-IMG5 | 51 | M | 50-59 | 30-39 | 45.3 | 42.1 | -8.9 | 0.814 | 36,216 |
| C36-IMG1 | 54 | F | 50-59 | 60-69 | 51.3 | 48.5 | -5.5 | 0.719 | 62,141 |
| C36-IMG4 | 54 | F | 50-59 | 70-79 | 56.0 | 54.4 | +0.4 | 0.543 | 63,006 |
| C36-IMG5 | 54 | F | 50-59 | 30-39 | 47.2 | 45.4 | -8.6 | 0.923 | 64,369 |
| C35-IMG2 | 64 | F | 60-69 | 60-69 | 54.3 | 54.3 | -9.7 | 0.909 | 48,021 |
| C35-IMG3 | 64 | F | 60-69 | 70-79 | 54.9 | 54.7 | -9.3 | 0.847 | 50,416 |
| C35-IMG4 | 64 | F | 60-69 | 70-79 | 51.2 | 52.2 | -11.8 | 0.954 | 52,400 |
| Total (24) | | | | | | | MAE=5.9 | | 2,399,803 |

**C**

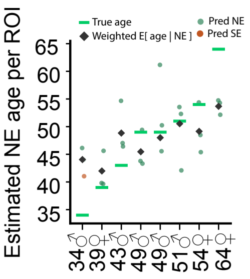


**D**

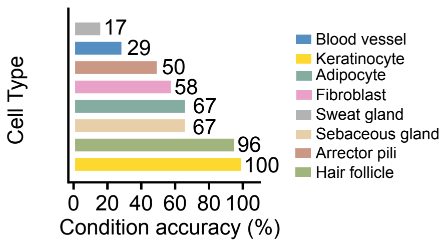
